# Distinct functions of tissue-resident and circulating memory Th2 cells in allergic airway disease

**DOI:** 10.1101/2020.03.25.006858

**Authors:** Rod A. Rahimi, Keshav Nepal, Murat Cetinbas, Ruslan I. Sadreyev, Andrew D. Luster

## Abstract

Memory CD4^+^ T helper type 2 (Th2) cells are critical in driving allergic asthma pathogenesis, yet the mechanisms whereby tissue-resident memory Th2 cells (Th2 Trm) and circulating memory Th2 cells collaborate *in vivo* remain unclear. Here, using a house dust mite (HDM) model of allergic asthma and parabiosis, we demonstrate that Th2 Trm and circulating memory Th2 cells perform non-redundant functions *in vivo*. Upon HDM re-challenge, circulating memory Th2 cells trafficked into the lung parenchyma and ignited perivascular inflammation to promote eosinophil and CD4^+^ T cell recruitment. In contrast, Th2 Trm proliferated near airways and promoted mucus metaplasia, airway hyper-responsiveness, and airway eosinophil activation. Transcriptional analysis revealed that Th2 Trm and circulating memory Th2 cells share a core Th2 gene signature, but also exhibit distinct transcriptional profiles. Specifically, Th2 Trm express a tissue adaptation signature, including genes involved in regulating and interacting with extracellular matrix. Our findings demonstrate that Th2 Trm and circulating memory Th2 cells are functionally and transcriptionally distinct subsets with unique roles in promoting allergic airway disease.

**SUMMARY:** How memory Th2 cell subsets orchestrate allergic airway inflammation remains unclear. Rahimi *et al*. use a murine model of allergic asthma and parabiosis to demonstrate that tissue-resident and circulating memory Th2 cells are functionally distinct subsets with unique roles in promoting allergic airway disease.

## Introduction

Asthma is an inflammatory airway disorder affecting more than 350 million individuals worldwide (Soriano et al., 2017). Although asthma is a heterogeneous syndrome, allergic airway inflammation drives asthma pathogenesis in the majority of children and half of adults (Arbes et al., 2007; Lambrecht and Hammad, 2015; Woodruff et al., 2009). The development of CD4^+^ T helper type 2 (Th2) cells that recognize airborne allergens is a key feature of allergic asthma (Lambrecht et al., 2019; Walker and McKenzie, 2018). Allergen-specific Th2 cells orchestrate allergic airway inflammation by producing type 2 cytokines, including IL-4, IL-5, and IL-13, which drive eosinophilic inflammation, mucus metaplasia, and airway hyper-responsiveness (Lambrecht et al., 2019; Walker and McKenzie, 2018). In addition, Th2 cells can give rise to long-lived memory Th2 cells that maintain allergen-specific immunity (Hondowicz et al., 2016; Onodera et al., 2018). Consequently, memory Th2 cells represent an attractive therapeutic target in allergic asthma, but our knowledge of memory Th2 biology *in vivo* remains limited.

Over the last twenty years, distinct subsets of memory T cells have been characterized that exhibit unique trafficking patterns and functions *in vivo* (Jameson and Masopust, 2018). Tissue-resident memory T cells (Trm) persist in previously inflamed non-lymphoid tissue (NLT), providing enhanced local immune memory (Carbone, 2015; Schenkel and Masopust, 2014). In contrast, circulating memory T cells provide global host defense (Jameson and Masopust, 2018). Most of our knowledge of Trm biology stems from the CD8^+^ T cell field with less known about CD4^+^ Trm. Parabiosis experiments have demonstrated that CD4^+^ T helper type 1 cells (Th1 Trm) are the dominant memory Th1 subset surveying NLT and initiating local recall responses (Beura et al., 2019). Both Th1 Trm and a “second wave” of recruited Th1 cells are required for optimal pathogen control *in vivo* (Stary et al., 2015; Iijima and Iwasaki, 2014; Glennie et al., 2015). Studies using the house dust mite (HDM) model of allergic asthma have shown that Th2 Trm persist in the lung in inducible bronchus-associated lymphoid tissue (iBALT) structures (Hondowicz et al., 2016; Shinoda et al., 2016; Turner et al., 2018). Interestingly, Th2 Trm can promote airway hyper-responsiveness and inflammatory cell recruitment even after depletion of circulating T cells, suggesting Th2 Trm are an important cell population orchestrating local type 2 immunity (Hondowicz et al., 2016; Turner et al., 2018). However, adoptively transferring circulating memory Th2 cells into naïve mice and administering repetitive antigen challenge leads to allergic airway inflammation (Endo et al., 2011, 2015). As a result, the mechanisms whereby Th2 Trm and circulating memory Th2 cells collaborate in an endogenous recall response are unknown, a gap in knowledge that limits therapeutic targeting of pathogenic memory Th2 cells in allergic airway disease.

Here, we use a HDM model of allergic asthma and parabiosis to define the function of endogenous tissue-resident and circulating memory Th2 cells. Unexpectedly, we found Th2 Trm and circulating memory Th2 cells performed distinct functions *in vivo*. Upon HDM re-challenge, circulating memory Th2 cells trafficked into the lung parenchyma and ignited perivascular inflammation to promote eosinophil and CD4^+^ T cell recruitment. In contrast, Th2 Trm proliferated near airways and promoted mucus metaplasia, airway hyper-responsiveness, and airway eosinophil activation. Transcriptional analysis revealed that Th2 Trm and circulating memory Th2 cells share a core Th2 gene signature, but also exhibit distinct transcriptional profiles. Specifically, Th2 Trm express a tissue adaptation signature, including genes involved in regulating and interacting with extracellular matrix. Our findings demonstrate that Th2 Trm and circulating memory Th2 cells are functionally and transcriptionally distinct subsets with unique roles *in vivo* with the establishment of Th2 Trm being critical for the full manifestation of allergic airway disease. We propose a novel model for memory Th2 responses in the airways with implications for developing disease-modifying therapies for individuals with allergic asthma.

## Results and Discussion

### Memory Th2 cells orchestrate the recall response to HDM in an allergen-specific manner

To define the function of endogenous memory Th2 cell subsets *in vivo*, we used a well-established murine model of allergic asthma via administration of intranasal (i.n.) HDM. HDM sensitization and repetitive challenge induces robust allergic airway inflammation that is dependent on Th2 cells and independent of IgE (Li et al., 2016; Hondowicz et al., 2016; McKnight et al., 2017). We sensitized and challenged mice with HDM followed by a 6-12 week rest period to generate “HDM-memory” mice (Fig. S1A). First, we assessed the ability of HDM-memory mice to exhibit a recall response *in vivo* by re-challenging with a single i.n. dose of HDM. Compared to naïve mice treated with a single dose of HDM, HDM-memory mice exhibited robust induction of type 2 cytokines and chemokines within the lung as well as the cardinal features of allergic inflammation, including lung eosinophilia, mucus metaplasia, and airway hyper-responsiveness (Fig. 1A-E and S1B-D). The observation that naïve mice do not exhibit a significant innate type 2 response after one dose of HDM suggests that memory Th2 cells are driving the recall response in HDM-memory mice. However, previous studies have demonstrated that allergen-exposure can lead to both memory Th2 cells as well as memory-like ILC2 with enhanced responsiveness to allergens (Martinez-Gonzalez et al., 2016, 2017). We sought to determine the relative contribution of memory Th2 cells and memory-like ILC2 to the recall response in HDM-memory mice. To do so, we initially assessed the proliferative response of memory Th2 cells and ILC2 to HDM re-challenge in HDM-memory mice. To examine Th2 cells in the lung parenchyma, we performed intravenous injection with fluorophore-labelled anti-CD45 antibody to label intravascular leukocytes prior to lung harvest as previously described (Fig. S2A) (Anderson et al., 2014; Galkina et al., 2005). We performed intra-nuclear staining for FoxP3 and GATA3, allowing us to identify Th2 cells as FoxP3^-^GATA3^+^CD4^+^ T cells (Fig. S2A). Consistent with previous reports, compared to naïve controls, we observed an increase in the number of Th2 cells persisting in the lung parenchyma of HDM-memory mice (Fig. 1F and S2B) (Hondowicz et al., 2016; Turner et al., 2018). Memory Th2 cells persisting in the lung parenchyma of HDM-memory mice expressed high levels of CD69, consistent with being Trm (Fig. S2C). Upon HDM treatment, naïve mice did not develop a significant increase in lung Th2 cells whereas HDM-memory mice exhibited a robust expansion of Th2 cells within the lung parenchyma and mediastinal lymph node (mLN) (Fig. 1F and G). In contrast, we did not observe a difference in the number of ILC2 between naïve and HDM-memory mice during homeostasis (Fig. 1H). In addition, ILC2 from HDM-memory mice minimally expanded upon HDM re-challenge (Fig. 1H).

**Figure 1.**
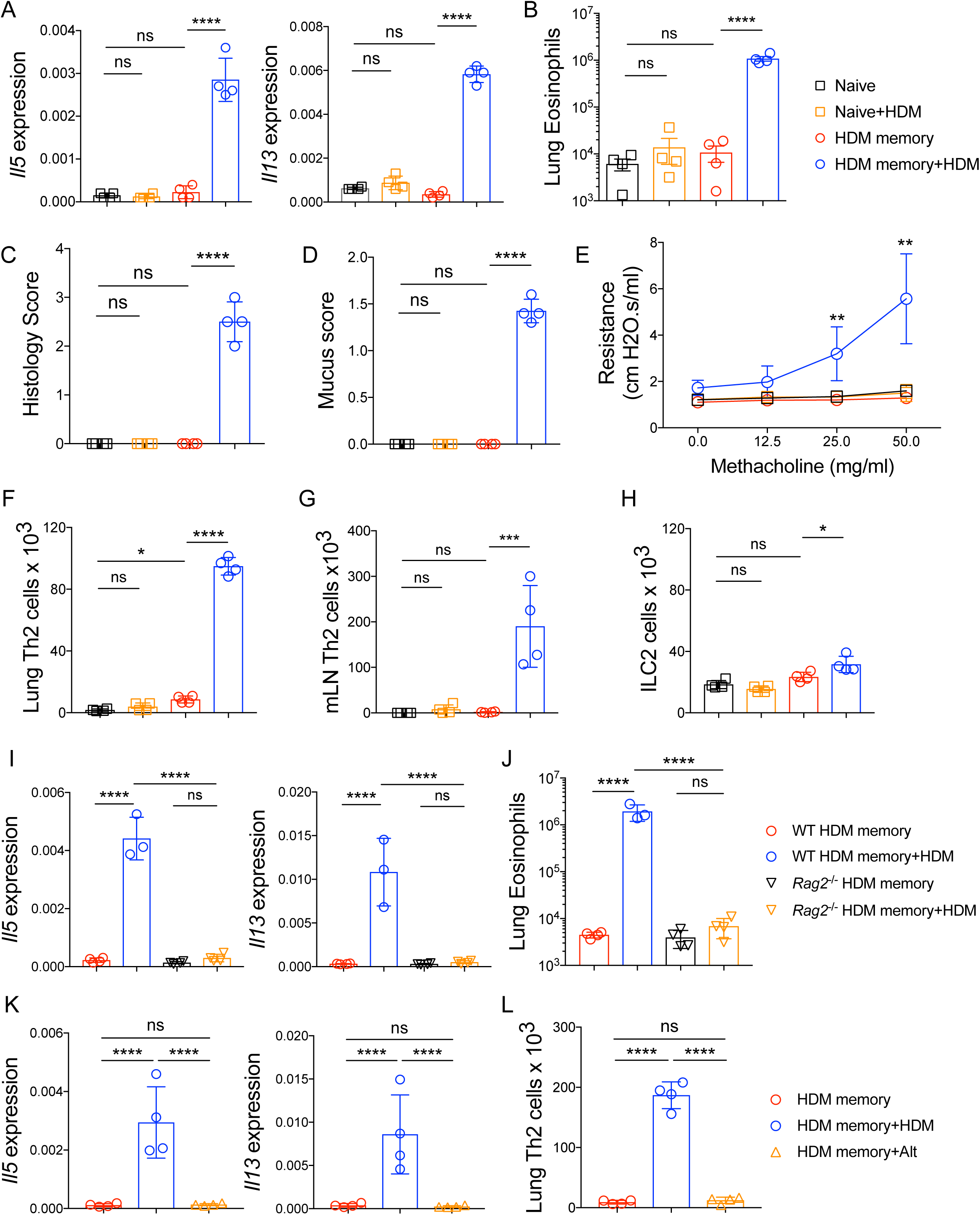
Memory Th2 cells orchestrate the recall response to HDM in an allergen-specific manner. (A-H) C57BL/6 mice were sensitized and challenged with intranasal (i.n.) HDM, rested for 6-12 weeks, and left untreated or re-challenged with a single dose of i.n. HDM followed by tissue harvest 72 hours later. (A) Lung *Il5* and *Il13* relative RNA levels assessed via qPCR. (B) Lung parenchymal (anti-CD45 i.v. unlabelled) eosinophils were quantitated via flow cytometry. (C) Lung histology scores. (D) Mucus scores. (E) Airway resistance was measured in indicated groups after increasing doses of methacholine. (F-G) Th2 cells (FoxP3^-^GATA3^+^CD4^+^ T cells) were quantitated by flow cytometry in the (F) lung parenchyma (anti-CD45 i.v. unlabeled) and (G) mLN in indicated groups. (H) Quantitation of lung intraparenchymal ILC2 (anti-CD45 i.v. unlabeled, lineage^-^CD4^-^Thy1.2^+^CD127^+^ST2^+^ cells). (I-J) C57BL/6 and *Rag2^-/-^* mice were sensitized and challenged with intranasal (i.n.) HDM, rested for 6-12 weeks, and left untreated or re-challenged with a single dose of i.n. HDM followed by tissue harvest 72 hours later. (I) Lung *Il5* and *Il13* relative RNA levels assessed via qPCR. (J) Lung parenchymal (anti-CD45 i.v. unlabeled) eosinophils were quantitated via flow cytometry. (K-L) C57BL/6 mice were sensitized and challenged with intranasal (i.n.) HDM, rested for 6-12 weeks, and left untreated or re-challenged with a single dose of i.n. HDM or i.n. *Alternaria* followed by tissue harvest 72 hours later. (K) Lung *Il5* and *Il13* relative RNA levels assessed via qPCR. (L) Th2 cells (FoxP3^-^GATA3^+^CD4^+^ T cells) were quantitated by flow cytometry in the lung parenchyma (anti-CD45 i.v. unlabeled). Representative data show individual mice with mean ± SEM from one of three independent experiments with 4 mice per group (A-H) or mean ± SEM from one of two independent experiments with 3-4 mice per group (I-L). One-way ANOVA analysis with Holm-Sidak’s testing was used for statistical analysis of multiple groups. *P < 0.05; **P < 0.01; ***P < 0.001.

To further investigate the potential role of memory-like ILC2 to the HDM recall response, we took advantage of the observation that allergen-experienced ILC2 cells can acquire memory-like properties independently of T cells (Martinez-Gonzalez et al., 2016). We generated wild-type (WT) and *Rag2^-/-^* HDM-memory mice and left them untreated or re-challenged with HDM. We found *Rag2^-/-^* HDM-memory mice failed to induce significant expression of type 2 cytokines or eosinophilia within the lung (Fig. 1I and J). These results indicate that HDM treatment promotes memory Th2 cell development, but does not efficiently induce memory-like ILC2 cells. These findings are consistent with previous studies demonstrating that Th2 cells are the functionally dominant type 2 lymphocyte in the primary response to HDM (Li et al., 2016; Hondowicz et al., 2016). Finally, given that Th2 cells are capable of responding to innate signals, including IL-33, in an antigen-independent manner, we investigated whether the memory Th2 cell recall response was allergen-specific (Guo et al., 2009, 2015; Minutti et al., 2017). To do so, we used *Alternaria alternata* fungal extract, which is well characterized to promote IL-33 release and allergic airway inflammation (Snelgrove et al., 2014; Causton et al., 2018). Specifically, we administered a single 10 μg dose of either HDM or *Alternaria* extract to HDM-memory mice. While HDM re-challenge induced robust type 2 cytokine expression and Th2 cell expansion, we did not observe increased expression of type 2 cytokines or Th2 cell expansion with *Alternaria* treatment (Fig. 1K and L). As a control, we generated *Alternaria*-memory mice and left them untreated or re-challenged with a single dose of *Alternaria*. *Alternaria* re-challenge promoted robust Th2 cell expansion within the lung of *Alternaria*-memory mice (Fig. S2D). In summary, memory Th2 cells proliferate and orchestrate allergic inflammation in the HDM model of allergic airway disease in an allergen-specific manner.

### Th2 Trm and circulating memory Th2 cells collaborate to promote Th2 cell expansion and type 2 cytokine production within the lung

Next, we sought to define the functions of Th2 Trm and circulating memory Th2 cells. Studies on Trm in other experimental systems have suggested that Trm are more potent cytokine producers than their circulating counterparts (Strutt et al., 2018; Smolders et al., 2018; Hombrink et al., 2016; Oja et al., 2018). To assess the ability of Th2 Trm and circulating memory Th2 cells to produce type 2 cytokines upon re-activation, we isolated lung CD4^+^ T cells from HDM-memory mice after anti-CD45 i.v. injection, performed *ex vivo* treatment with anti-CD3 and anti-CD28, and assessed the ability of intraparenchymal Th2 Trm (anti-CD45 i.v. negative) and intravascular memory Th2 cells (anti-CD45 i.v. positive) to produce type 2 cytokines (Fig. S3A and S3B). While both memory Th2 cell subsets are capable of producing IL-5 and IL-13, we found Th2 Trm within the lung parenchyma produced a greater amount of type 2 cytokines on a per cell basis (Fig. S3A and S3B).

To investigate the mechanisms whereby Th2 Trm enhance the HDM recall response *in vivo*, we utilized a parabiosis system in which we surgically conjoined congenic HDM-memory mice (CD45.2 memory parabiont) and naïve mice (CD45.1 naïve parabiont). Parabiosis leads to a shared vasculature such that circulating memory Th2 cells equilibrate between both parabionts, but only the CD45.2 memory parabiont have Th2 Trm. After 3-4 weeks of parabiosis, we administered a single dose of i.n. HDM to both parabionts (Fig. 2A). First, we assessed whether the parabiosis system had effectively led to chimerism of the circulating memory T cell compartment. While CD45.1 naïve non-parabiotic mice administered a single dose of HDM did not induce an increase in Th2 cells within the mLN after 72 hours, naïve and memory parabionts exhibited a robust population of Th2 cells in the mLN (Fig. 2B). To confirm that the Th2 cells within the mLN of both parabionts were derived from memory Th2 cells, we assessed the expression of CD45.2 within the GATA3^-^ and GATA3^+^CD4^+^ T cell populations. GATA3^+^CD4^+^ T cells from the mLN were overwhelming derived from the CD45.2 memory parabiont, whereas GATA3^-^CD4^+^ T cells showed ∼50:50 chimerism (Fig. 2C). In addition, quantitation of Th2 cells from the mLN revealed similar numbers of Th2 cells in the two parabionts (Fig. 2D). Finally, total mLN cells from the naïve and memory parabionts re-stimulated *ex vivo* with HDM produced similar levels of IL-5 and IL-13 (Fig. 2E). In summary, these results demonstrate that parabiosis effectively transferred endogenous, circulating memory Th2 cells *in vivo*.

**Figure 2.**
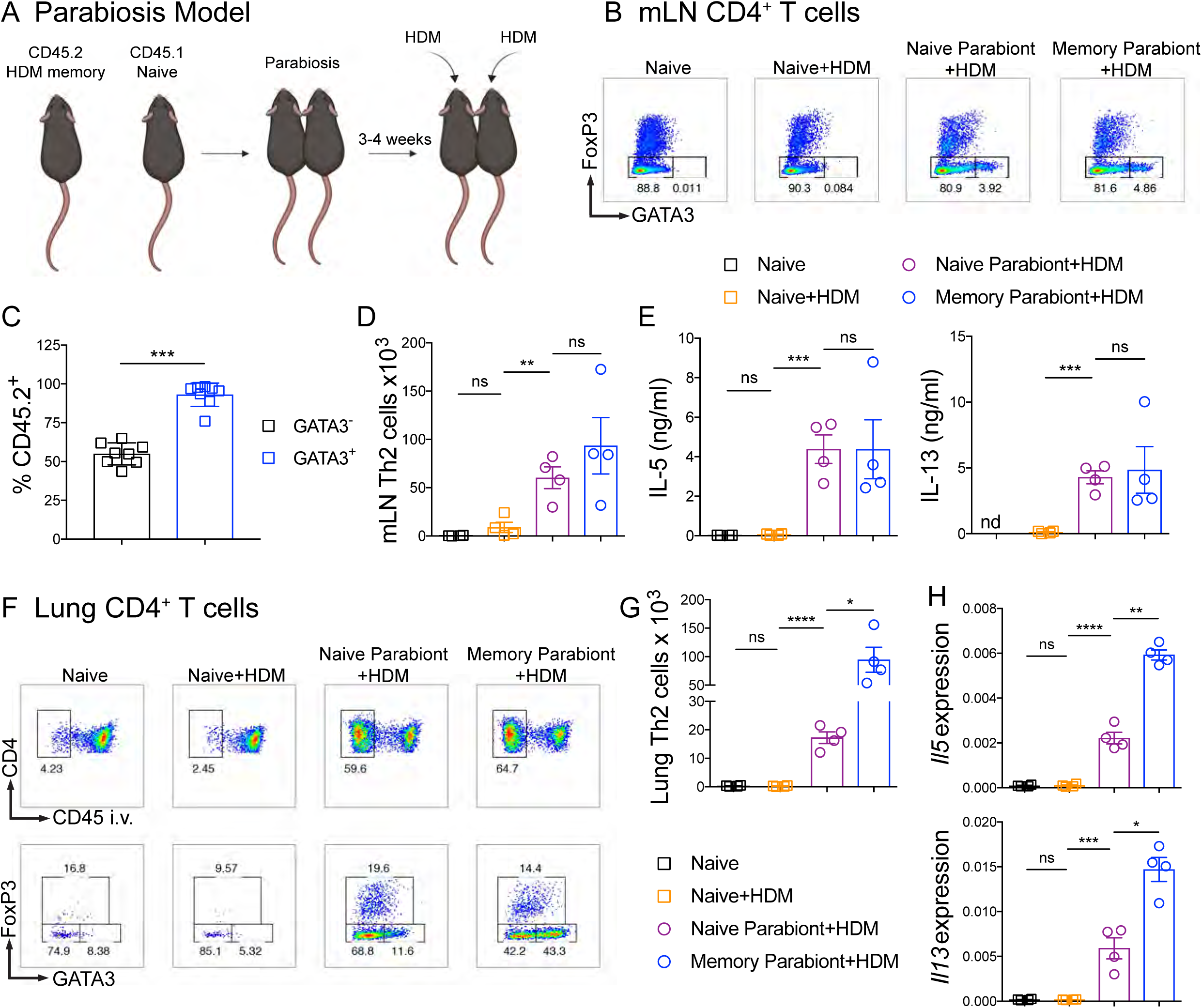
Th2 Trm and circulating memory Th2 cells collaborate to promote Th2 cell expansion and type 2 cytokine production within the lung. (A-H) CD45.2 HDM-memory mice were surgically conjoined to CD45.1 naïve mice. After 3-4 weeks both parabionts received a single dose of i.n. HDM with harvest of mLN and lung after 72 hours. (A) Schematic of parabiosis experiment. (B) Representative flow cytometry of mLN CD4^+^ T cells isolated from indicated groups demonstrating FoxP3 and GATA3 expression. (C) Percentage of CD45.2^+^ cells (from memory parabiont) of GATA3^-^ and GATA3^+^ CD4^+^ T cells from the mLN of naïve and memory parabionts. (D) Quantitation of Th2 cells from mLN from indicated groups. (E) Total mLN cells were re-stimulated *ex vivo* with 25 μg/ml HDM for 72 hours and levels of IL-5 and IL-13 protein in the supernatant was measured via ELISA. (F) Representative flow cytometry of lung CD4^+^ T cells from indicated groups demonstrating FoxP3 and GATA3 expression. (G) Lung Th2 cells (anti-CD45 i.v. unlabeled, FoxP3^-^GATA3^+^CD4^+^ T cells) were quantitated via flow cytometry. (H) Lung *Il5* and *Il13* relative RNA levels assessed via qPCR. Representative data show individual mice with mean ± SEM from one of three independent experiments with 3-4 mice per group. For statistical analysis, one-way ANOVA analysis with Holm-Sidak’s testing was used for statistical analysis of multiple groups with paired two-tailed t tests for comparison on naive and memory parabiont groups. *P < 0.05; **P < 0.01; ***P < 0.001.

Next, we turned to the HDM recall response within the lung. Within the lung parenchyma (anti-CD45 i.v. negative), CD45.1 naïve, non-parabiotic mice had few CD4^+^ T cells and after a single dose of HDM there was no increase (Fig. 2F and G). In contrast, naïve parabionts possessing circulating memory Th2 cells exhibited a robust accumulation of CD4^+^ T cells within the lung parenchyma (Fig. 2F and G). However, memory parabionts had significantly more Th2 cells (defined by being FoxP3^-^GATA3^+^CD4^+^ cells) within the lung (Fig. 2F and G). In support of Th2 Trm driving an enhanced type 2 response, expression of type 2 cytokines within the lung was higher in memory parabionts than naïve parabionts following HDM challenge (Fig. 2H). Of note, naïve parabionts did induce greater type 2 cytokine expression compared to naïve, non-parabiotic mice treated with HDM. Consequently, we found that Th2 Trm and circulating memory Th2 cells collaborate to promote Th2 cell expansion and type 2 cytokine production within the lung upon HDM re-challenge.

### Th2 Trm and circulating memory Th2 cells perform non-redundant functions upon HDM re-challenge

To define the roles of Th2 Trm and circulating memory Th2 cells in promoting the cardinal features of allergic airway inflammation within the lung, we first enumerated the number of eosinophils within the lung parenchyma of naïve and memory parabionts as well as naïve controls after HDM challenge. Unexpectedly, we found comparable numbers of eosinophils within the lung parenchyma and bronchoalveolar lavage (BAL) of both parabionts, suggesting that circulating memory Th2 cells were necessary and sufficient to promote eosinophil mobilization and recruitment into the lung and airways (Fig. 3A, Fig. S3C and S3D). The eosinophil-attracting chemokines CCL11 (eotaxin-1) and CCL24 (eotaxin-2) are induced by IL-4 and IL-13 and are well characterized to promote eosinophil recruitment into the lung parenchyma via CCR3 (Griffith et al., 2014). Consequently, we compared *Ccl11* and *Ccl24* expression within the lung of naïve and memory parabionts as well as naïve controls after HDM challenge. Compared to naïve controls, naïve parabionts acquired the ability to induce expression of *Ccl11* and *Ccl24*, but to levels less than memory parabionts (Fig. 3B). Consequently, it remained unclear why the enhanced CCR3 ligand expression within the lungs of the memory parabionts did not result in greater eosinophil recruitment into the lung. Since CCL11 and CCL24, as well as IL-5, have been shown to induce recruitment as well as activation of eosinophils, we investigated the expression of several well-characterized eosinophil activation markers, including CD11b and CD62L, which are upregulated and downregulated, respectively, upon eosinophil activation (Johansson, 2014; Lukacs, 2001). We found significantly higher expression of CD11b and lower expression of CD62L on eosinophils present in the BAL of memory parabionts, consistent with enhanced activation (Fig. 3C). These findings demonstrate that Th2 Trm and circulating memory Th2 cells both contribute to *Ccl11* and *Ccl24* expression within the lung upon HDM re-challenge. In addition, our results suggest that Th2 Trm-driven CCL11 and CCL24 production does not significantly enhance the recruitment of eosinophils from the blood into the lung parenchyma; instead, tissue-resident memory enhanced eosinophil activation within the airways.

**Figure 3.**
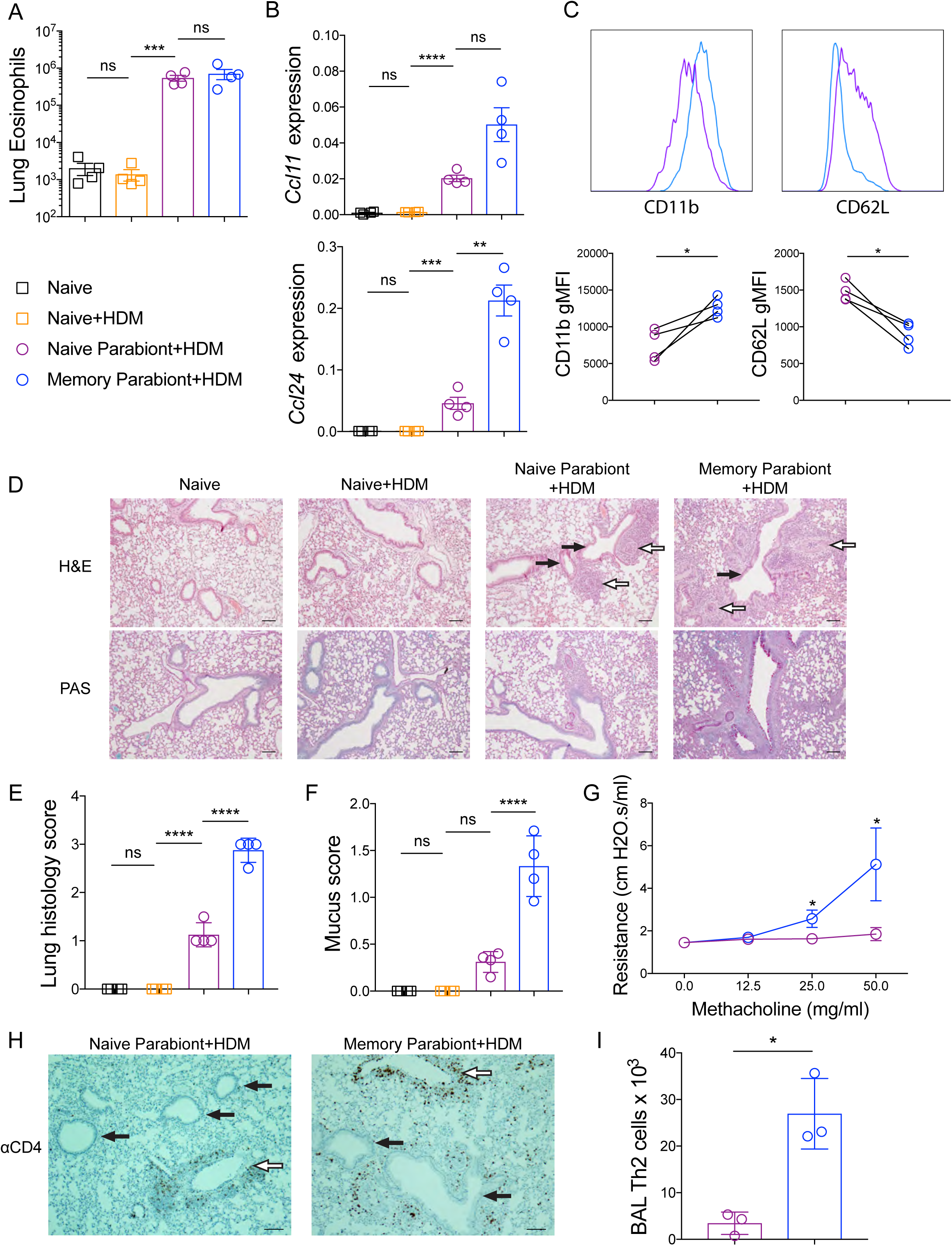
Th2 Trm and circulating memory Th2 cells perform non-redundant functions upon HDM re-challenge. (A) Quantitation of eosinophils (anti-CD45 i.v. unlabeled, CD11c^lo^Siglec-F^+^ cells) from the lung of indicated groups. (B) Lung *Ccl11* and *Ccl24* relative RNA levels assessed via qPCR. (C) Representative histograms demonstrating BAL eosinophil cell surface expression of CD11b and CD62L (top panel) and geometric mean fluorescence intensity (gMFI) (bottom panel) from naïve parabionts (purple circles) and memory parabionts (blue circles) as determined by flow cytometry. (D) H&E-stained (top panel) and PAS-stained (bottom panel) lung sections from indicated groups. White arrows indicate blood vessels, black arrows indicate airways. (E) Lung histology scores. (F) Mucus scores. (G) Airway resistance was measured in indicated group after increasing doses of methacholine. (H) Immunohistochemistry for lung CD4^+^ T cells in naïve and memory parabionts 72 hours after HDM challenged. White arrows indicate blood vessels, black arrows indicate airways. (I) BAL Th2 cells (FoxP3^-^GATA3^+^CD4^+^ T cells) were quantitated in indicated groups via flow cytometry. Representative data show individual mice with mean ± SEM from one of three independent experiments with 3-4 mice per group (A-F, H-I) or mean ± SEM from two cumulative experiments with 4 mice per group (G). For statistical analysis, one-way ANOVA analysis with Holm-Sidak’s testing was used for statistical analysis of multiple groups with paired two-tailed t tests for comparison on naive and memory parabiont groups. *P < 0.05; **P < 0.01; ***P < 0.001.

We then assessed the pattern and severity of inflammation within the lung parenchyma from naïve and memory parabionts as well as naïve controls after HDM treatment. The lung inflammation induced by circulating memory Th2 cells in the naïve parabiont mice was mainly perivascular, consistent with our data that recruited Th2 cells are sufficient for eosinophil recruitment (Fig. 3D). In contrast, memory parabionts with Th2 Trm exhibited more peribronchial inflammation than naïve parabionts (Fig. 3D). Memory parabionts had higher lung inflammation scores, predominantly due to airway inflammation (Fig. 3E). In support of the observation that Th2 Trm were required for efficient peribronchial inflammation, we found greater mucus metaplasia in memory parabionts (Fig. 3D and 3F). In murine models of allergic asthma, allergen treatment and administration of methacholine increases airway resistance, which requires airway mucus production (Evans et al., 2015). To assess the role of Th2 Trm and circulating memory Th2 cells in airway hyper-responsiveness, we measured airway resistance in naïve and memory parabionts after HDM treatment and increasing doses of methacholine. In support of our histologic findings, increased airway resistance only occurred in memory parabionts re-challenged with HDM (Fig. 3G). We also found that CD4^+^ T cells in naïve parabionts treated with HDM localized around blood vessels, and did not accumulate near airways, whereas memory parabionts treated with HDM exhibited accumulations of CD4^+^ T cells around both blood vessels and airways (Fig. 3H). Finally, we performed BAL in naïve and memory parabionts challenged with HDM followed by flow cytometry for FoxP3^-^GATA3^+^CD4^+^ T cells and found markedly greater numbers of Th2 cells in the airways of memory parabionts (Fig. 3I). These results demonstrate that Th2 Trm and circulating memory Th2 cells both contribute to the recall response to HDM and perform non-redundant function *in vivo*. Circulating memory Th2 cells are sufficient to promote perivascular inflammation and eosinophil recruitment, while Th2 Trm promote peribronchial inflammation, including mucus metaplasia, airway hyper-responsiveness, and airway eosinophil activation.

### Th2 Trm drive the airway Th2 response to HDM via *in situ* proliferation, but are dispensable for CD4^+^ T cell recruitment

We subsequently turned to the mechanisms regulating the trafficking of Th2 Trm and circulating memory Th2 cells during homeostasis and HDM re-challenge. Although Trm are the dominant memory T cell population in non-lymphoid tissue (NLT) during homeostasis, a population of re-circulating (non-resident) memory T cells can dynamically survey NLT (Bromley et al., 2013; Gerlach et al., 2016). In addition, in a murine model of influenza infection, lung CD8^+^ Trm wane with time due to apoptosis with a half-life of 5 days and require circulating memory T cells to “re-seed” the Trm compartment during homeostasis (Slütter et al., 2017). To investigate the extent to which circulating memory Th2 cells re-circulate or re-seed the Th2 Trm pool within the lung parenchyma during homeostasis, we surgically conjoined congenic (CD45.2 and CD45.1) HDM-memory mice (Fig. 4A). After 3-4 weeks, we found that memory Th2 cells within the lung parenchyma were overwhelming derived from the host, indicating the absence of significant memory Th2 cell re-circulation or circulating memory Th2 cells “re-seeding” the Th2 Trm pool in the HDM model, at least during this time frame (Fig. 4B and C). In contrast, FoxP3^-^GATA3^-^CD4^+^ T cells (non-Th2 cells) within the lung parenchyma were a mix of tissue-resident and re-circulating memory CD4^+^ T cells (Fig S3E). To investigate Th2 cell trafficking during HDM re-challenge, we administered a single dose of HDM to both memory parabionts and found a significant influx of partner-derived Th2 cells (Fig. 4C). Consequently, Th2 Trm do not require re-seeding from circulating memory Th2 cells and circulating memory Th2 cells only traffic into the lung parenchyma upon HDM re-challenge.

**Figure 4.**
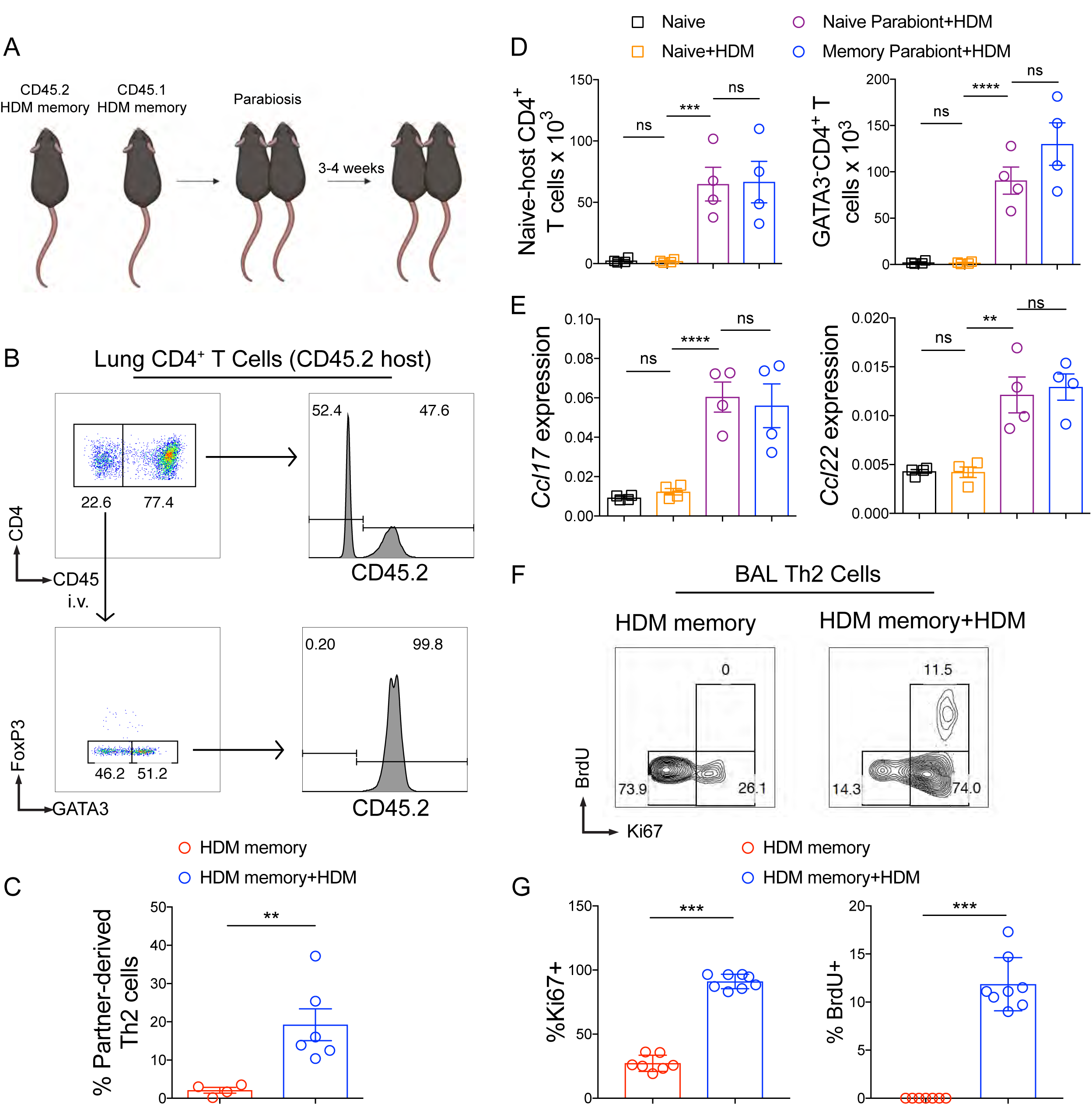
Th2 Trm drive the airway Th2 response to HDM via *in situ* proliferation, but are dispensable for CD4^+^ T cell recruitment. (A-B) CD45.2 and CD45.1 HDM-memory mice were surgically conjoined. After 3-4 weeks both HDM-memory parabionts were left untreated or received a single dose of i.n. HDM with harvest of lung after 72 hours. (A) Schematic of parabiosis experiment. (B) Representative flow cytometry of lung CD4^+^ T cells from untreated CD45.2 memory parabiont demonstrating CD45.2 expression of anti-CD45 labelled CD4^+^ T cells and FoxP3 and GATA3 expression of anti-CD45 unlabelled CD4^+^ T cells as well as CD45.2 expression of FoxP3^-^GATA3^+^CD4^+^ T cells. (C) Quantitation of percent partner-derived lung Th2 cells (anti-CD45 i.v. unlabeled, FoxP3^-^GATA3^+^CD4^+^ T cells) assessed via flow cytometry from untreated and HDM re-challenged memory parabionts. (D-E) CD45.2 HDM-memory mice were surgically conjoined to CD45.1 naïve mice. After 3-4 weeks both parabionts received a single dose of i.n. HDM with harvest of lung after 72 hours. (D) Lung naïve-host-derived CD4^+^ T cells (anti-CD45 i.v. unlabeled, FoxP3^-^CD45.2^-^CD4^+^ T cells) and anti-CD45 i.v. unlabeled FoxP3^-^GATA3^-^CD4^+^ T cells were quantitated via flow cytometry. (E) Lung *Ccl17* and *Ccl22* relative expression assessed via qPCR. (F-G) Individual HDM-memory mice were left untreated or received a single dose of i.n. HDM with BAL after 72 hours. BrdU was injected i.p. 2 hours prior to BAL. (F) Representative flow cytometry of BAL Th2 cells (FoxP3^-^GATA3^+^CD4^+^ T cells) from HDM-memory mice without and with HDM re-challenge demonstrating Ki67 and BrdU expression. (G) Quantitation of percent Ki67^+^ and BrdU^+^ BAL Th2 cells in indicated groups. Representative data show individual mice with mean ± SEM from two cumulative experiments with 4-6 mice per group (A-C) or mean ± SEM from one of three independent experiments with 3-4 mice per group (D-E) or mean ± SEM from two cumulative experiments with 8 mice per group (F-G). For statistical analysis, two-tailed Mann-Whitney U testing was performed for non-parametric data. One-way ANOVA analysis with Holm-Sidak’s testing was used for statistical analysis of multiple groups with paired two-tailed t tests for comparison on naive and memory parabiont groups. *P < 0.05; **P < 0.01; ***P < 0.001.

Next, we investigated the mechanism whereby tissue-resident memory to HDM enhances Th2 cell expansion (as demonstrated in Fig. 2F and G). Trm within NLT have been demonstrated to enhance local expansion of T cells in two ways. First, in models of type 1 immunity, CD8^+^ and Th1 Trm have been shown to enhance the recruitment of circulating T cells (Schenkel et al., 2013; Glennie et al., 2015). Second, while Trm have long been assumed to be terminally-differentiated and proliferate poorly, recent studies have shown that CD8^+^ Trm can proliferate *in situ* (Park et al., 2018; Beura et al., 2018). To determine whether Th2 Trm promote enhanced CD4^+^ T cell recruitment from the circulation, we returned to our parabiosis system involving HDM-memory and naïve mice. Specifically, to measure CD4^+^ T cell recruitment, we investigated the number of conventional CD4^+^ T cells derived from the naïve mouse that trafficked into the lung parenchyma upon HDM challenge. CD45.1 naïve, non-parabiotic mice treated with HDM did not exhibit CD4^+^ T cell recruitment, suggesting innate signals induced by a single dose of HDM are insufficient to promote CD4^+^ T cell recruitment (Fig. 4D). In contrast, both naïve and memory parabionts exhibited similar recruitment of naïve-host derived CD45.2^-^CD4^+^ T cells into the lung parenchyma (Fig. 4D). In addition, the number of FoxP3^-^GATA3^-^CD4^+^ T cells (non-Th2 cells) and Tregs in the lung parenchyma were similar in the two parabionts, suggesting that tissue-resident memory is dispensable for CD4^+^ T cell recruitment (Fig. 4D and S3F). The chemokine receptor CCR4 is well-described to promote CD4^+^ T cell recruitment into the lung during type 2 immune responses via expression of the CCR4 ligands CCL17 and CCL22, which are induced by IL-4 and IL-13 (Mikhak et al., 2009, 2013; Perros et al., 2009; Faustino et al., 2013; Afshar et al., 2013). We examined the expression of *Ccl17* and *Ccl22* after HDM administration in naïve and memory parabionts as well as naïve controls. CD45.1 naïve, non-parabiotic mice treated with HDM did not exhibit increased expression of either chemokine in the lung after HDM administration. In contrast, naïve parabionts treated with HDM induced *Ccl17* and *Ccl22* expression in the lung to similar levels as memory parabionts, consistent with our observations of similar Treg and CD4^+^ T cell recruitment (Fig. 4E). Consequently, in agreement with circulating memory Th2 cells being sufficient to promote perivascular inflammation and lung eosinophilia, we found circulating memory Th2 cells were sufficient to promote CD4^+^ T cell recruitment into the lung parenchyma upon HDM treatment.

The observations that Th2 Trm enhance the number of Th2 cells within the lung and BAL without increasing CD4^+^ T cell recruitment suggested that Th2 Trm enhance the Th2 response within the airways primarily by proliferating *in situ*. To confirm this, we took advantage of our observation that Th2 Trm are required for Th2 cell expansion in the BAL (demonstrated in Fig. 3I). We left individual HDM-memory mice untreated or re-challenged with a single dose of HDM. Two hours prior to harvest, we administered an intraperitoneal injection of BrdU and performed BAL to assess Ki67 expression and BrdU uptake in BAL Th2 cells. During homeostasis, the majority of BAL Th2 cells exhibited low expression of Ki67 and did not uptake BrdU, consistent with being in the G0 phase of the cell cycle (Fig. 4F and G). Upon HDM re-challenge, airway Th2 cells dramatically increased in number and increased Ki67 expression with a subset acquiring BrdU (Fig. 4F, G and S3G). Our various experimental approaches demonstrate that Th2 Trm are the dominant memory Th2 cell subset surveying the lung parenchyma during homeostasis and respond to HDM re-challenge by robustly proliferating *in situ* near airways, but are dispensable for CD4^+^ T cell recruitment into the lung parenchyma.

### Shared and distinct transcriptional profiles of Th2 Trm and circulating memory Th2 cells

Studies on CD8^+^ Trm and Th1 Trm have demonstrated that these tissue-resident memory T cells exhibit a distinct transcriptional profile from their circulating counterparts (MacKay et al., 2013; Wakim et al., 2012; Pan et al., 2017; Mackay et al., 2016; Strutt et al., 2018; Beura et al., 2019). Given the distinct trafficking patterns and functions of Th2 Trm and circulating memory Th2 cells that we have elicited here, we sought to define the transcriptional profiles of these two memory Th2 cell subsets via RNA-seq analysis. We used FoxP3*^YFPCre^* reporter mice to generate HDM-memory mice and then sorted YFP^-^ ST2^+^CD4^+^ T cells from the lung parenchyma and secondary lymphoid organs (Fig. 5A). We took this approach to avoid fixation and permeabilization, which is required for intranuclear staining for FoxP3 (to exclude Tregs) and GATA3 (to identify Th2 cells). Rather, we used YFP fluorescence to exclude Tregs and the IL-33 receptor ST2 to identify Th2 cells, the latter marker having been shown to be a reliable surface marker for Th2 cells in the murine HDM model (Tibbitt et al., 2019). Along with the above memory Th2 cell populations, we also sorted YFP^-^ST2^-^CD4^+^ T cells from the lung and YFP^-^ST2^-^CD44^+^CD4^+^ T cells SLO as control populations of non-T helper type 2 cells. We excluded CD44^-^CD4^+^ T cells from the SLO to avoid a mixed population of naïve and memory CD4^+^ T cells in our control population. Principal component analysis of differentially expressed genes demonstrated that lung ST2^+^CD4^+^ T cells and SLO ST2^+^CD4^+^ T cells were transcriptionally distinct subsets (Fig. 5B). We next compared canonical T helper gene signatures in our various memory CD4^+^ T cell populations. As expected, Th1, Th17, and Tfh gene signatures were enriched in the memory ST2^-^CD4^+^ T cells from the lung and SLO (Fig. 5C). Lung and SLO memory ST2^+^CD4^+^ T cells exhibited similar expression of many canonical Th2 genes, including *Gata3*, *Il1rl1* (ST2), *Il17rb1* (IL-25R), *Il4*, *Il5*, and *Il3* (Fig. 5C). Lung memory ST2^+^CD4^+^ T cells expressed higher levels of the IL-13 receptor *Il13ra1* compared to SLO memory ST2^+^CD4^+^ T cells, indicating that Th2 Trm exhibit enhanced responsiveness to IL-13 within the lung parenchyma (Fig. 5C). A recent study performing single cell RNA-seq analysis on CD4^+^ T cells from the primary HDM model demonstrated that effector Th2 cells within the lung are a predominant source of IL-6 and also identified several Th2-specific transcripts, including *Pparg*, *Cd200r1*, *Igfbp7*, *Plac8* (Tibbitt et al., 2019). PPAR*γ* has been shown to be required for effective induction of IL-5 and IL-13 in effector Th2 cells (Angela et al., 2016; Chen et al., 2017; Nobs et al., 2017; Tibbitt et al., 2019). However, whether these Th2 cell-specific genes are expressed in the memory phase and the relative expression within Th2 Trm and circulating memory Th2 cells is not known. We found lung and SLO memory ST2^+^CD4^+^ T cells express similar levels of *Il6*, *Pparg*, and *Cd200r1* (Fig. 5C). In contrast, lung memory ST2^+^CD4^+^ T cells expressed higher levels of *Igfbp7* and *Plac8* than SLO memory ST2^+^CD4^+^ T cells (Fig. 5C). While the function of *Igfbp7* and *Plac8* in Th2 cell biology remains unclear, our results suggest these genes may play an important role in Th2 Trm function. Lastly, while the chemokine receptors CCR4 and CCR8 have been shown to be expressed by Th2 cells (Griffith et al., 2014), we found differential expression in lung and SLO memory ST2^+^CD4^+^ T cells with *Ccr4* being preferentially expressed in SLO memory ST2^+^CD4^+^ T cells and *Ccr8* being preferentially expressed in lung memory ST2^+^CD4^+^ T cells (Fig. 5C), indicating these two memory Th2 cell subsets differentially respond to type 2 chemokines *in vivo*.

**Figure 5.**
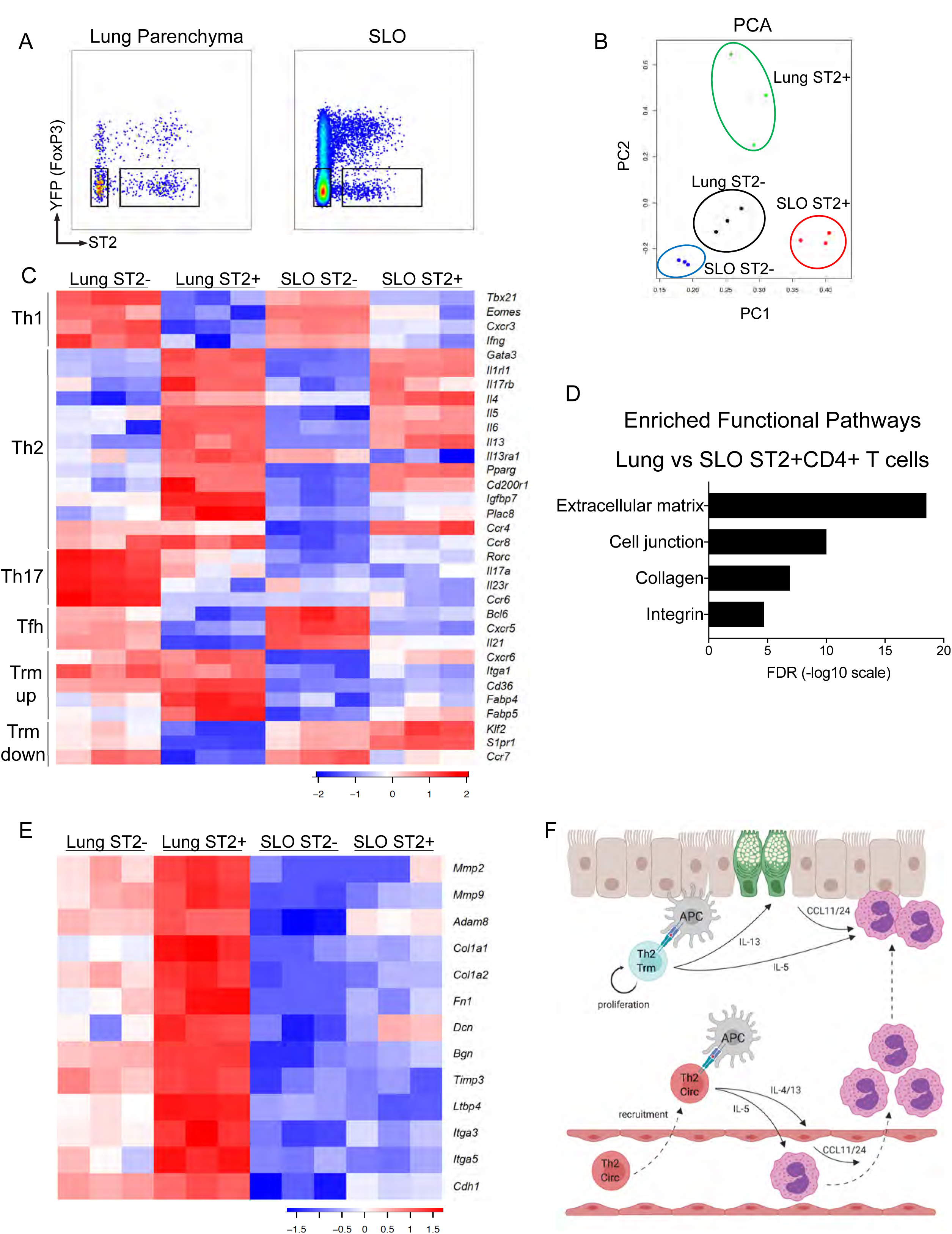
Shared and distinct transcriptional profiles of Th2 Trm and circulating memory Th2 cells. (A-E) FoxP3*^YFPCre^* mice were sensitized and challenged with intranasal HDM and rested for 6 weeks followed by lung and LN/spleen (SLO) harvest. (A) Representative flow cytometry of lung and SLO CD4^+^ T cells indicating YFP and ST2 expression as well as sorting gates for YFP^-^ST2^-^ and YFP^-^ST2^+^ populations. (B) Principal component Analysis analysis (PCA) plot based on differentially expressed genes from RNA-seq data of indicated memory CD4^+^ T cell populations. (C) Heatmap of expression values of T cell subset genes including Th1, Th2, Th17, Tfh, and Trm upregulated and downregulated genes. (D) Bar graph of statistical significance (false discovery rate (FDR) by the DAVID tool) of pathway enrichment among genes overexpressed in lung YFP^-^ST2^+^CD4^+^ T cells relative to SLO YFP^-^ST2^+^CD4^+^ T cells. (E) Heatmap of a selected subset of genes known to be involved in extracellular matrix biology and cell adhesion. Data are from three independent experiments with five HDM-memory mice pooled for each experiment (cell sort and RNA isolation). (F) Model of memory Th2 cell subsets during allergen re-challenge. Transcriptionally distinct memory Th2 cell subsets include tissue-resident memory Th2 cells persisting in the lung parenchyma (Th2 Trm indicated in blue) and circulating memory Th2 cells re-circulating in the blood (Th2 Circ indicated in red). During homeostasis, Th2 Trm and circulating memory Th2 cells exhibit non-overlapping trafficking patterns. Upon allergen re-exposure, circulating memory Th2 cells traffic into the lung parenchyma and produce type 2 cytokines upon antigen recognition by antigen presenting cells (APC), promoting perivascular inflammation and eosinophil recruitment. Activation of Th2 Trm leads to *in situ* proliferation and production of type 2 cytokines near the airways, which promotes mucus metaplasia (green epithelial cells) and enhances eosinophil activation.

Next, we assessed the expression of genes known to play a role in the maintenance and/or function of Trm *in vivo*. CD8^+^ Trm express the chemokine receptor CXCR6 and CD49a, which constitutes the *α*-subunit of the *α*1*β*1 integrin and binds to collagen (Szabo et al., 2019). CD8^+^ Trm within the lung require CXCR6 to localize to airways and α1β1 is required for effective CD8^+^ T cell immunity within the lung (Wein et al., 2019; Richter and Topham, 2007; Ray et al., 2004). We found Th2 Trm express higher levels of *Cxcr6* as well as CD49a (*Itga1*) than circulating memory ST2^+^CD4^+^ T cells (Fig. 5C). In addition, CD8^+^ Trm have been shown to require exogenous lipid uptake and metabolism, including fatty-acid binding proteins, for survival in NLT (Pan et al., 2017). Similar to CD8^+^ Trm, Th2 Trm express higher levels of the lipid scavenging molecule *Cd36* as well as fatty-acid binding proteins (*Fabp4* and *Fabp5*) than circulating memory Th2 cells (Fig. 5C) (Pan et al., 2017). Lastly, the establishment of CD8^+^ Trm, as well as tissue-resident NKT and NK cells, requires the transcription factors Blimp1 and Hobit, which directly promote the downregulation of pathways involved in egress from non-lymphoid tissues, including the genes *Klf2*, *S1pr1*, and *Ccr7* (Mackay et al., 2016). We found Th2 Trm downregulated tissue egress genes relative to circulating memory Th2 cells (Fig. 5C). However, we did not find that Th2 Trm preferentially express *Prdm1* (Blimp1) or *Znf683* (Hobit) or the CD8^+^ Trm-promoting transcription factor *Runx3* (Milner et al., 2017; Mackay et al., 2016). Transcription factors critical to Trm differentiation, such as Runx3, can be expressed at similar levels in Trm and circulating memory T cells, indicating that transcription factor function, rather than relative expression, drives Trm development (Milner et al., 2017).

In order to identify additional unique pathways differentially expressed between lung and SLO memory ST2^+^CD4^+^ T cells, we performed pathway enrichment analysis using the DAVID tool (Huang et al., 2009). This analysis of lung and SLO memory ST2^+^CD4^+^ T cells in our HDM model revealed the enrichment of extracellular matrix (ECM)-regulating and interacting genes among differentially expressed genes (Fig. 5D). Compared to SLO memory ST2^+^CD4^+^ T cells, lung memory ST2^+^CD4^+^ T cells expressed higher transcript levels of metalloproteases, such as *Mmp2*, *Mmp9* and *Adam8*, ECM components, such as collagen (*col1a1* and *col1a2*), fibronectin (*fn1*), Decorin (*Dcn*), Byglycan (*Bgn*), as well as regulators of ECM, including tissue inhibitor of metalloproteases *Timp3* and the TGF-*β* binding protein *Ltbp4* (Fig. 5E). In addition, genes involved in cell-to-ECM and cell-to-cell interaction, including *Itga3*, *Itga5*, and *Cdh1,* were enriched in lung memory ST2^+^CD4^+^ T cells compared to SLO memory ST2^+^CD4^+^ T cells (Fig. 5E). Our findings indicate that Th2 Trm and circulating memory Th2 cells share a core Th2 gene signature, while Th2 Trm uniquely express a tissue adaptive program, which includes genes involved in lipid metabolism and ECM biology. Notably, metalloproteases, ECM components, and specific integrins have also been shown to be enriched in CD8^+^ and Th1 Trm in other experimental systems, suggesting that regulation and interaction with the ECM may play an important role in the development, maintenance, and function of Trm in various tissues (MacKay et al., 2013; Wakim et al., 2012; Pan et al., 2017; Beura et al., 2019). Lastly, a recent study using Aspergillus fungal extract to promote airway inflammation in mice found that effector CD4^+^ T cells expressing CD69 are enriched for genes involved in ECM biology and fibrosis, indicating that effector CD4^+^ T cells are also capable of directly regulating the ECM within the lung during active inflammation (Ichikawa et al., 2019).

Previous studies investigating the function of Th2 Trm have relied on strategies to decrease the numbers of circulating T cells, including treatment with the S1P1 agonist FTY720 or anti-Thy1 antibody (Hondowicz et al., 2016; Turner et al., 2018; Ichikawa et al., 2019). Notably, decreasing circulating T cells during allergen re-challenge did not significantly reduce the number of allergen-specific CD4^+^ T cells within the lung or the allergen recall response, suggesting circulating memory Th2 cells do not play a significant role (Hondowicz et al., 2016; Turner et al., 2018; Ichikawa et al., 2019). While FTY720 efficiently sequesters naïve T cells in lymph nodes, it is less efficient at causing lymphopenia of memory T cells (Hofmann et al., 2006). In addition, FTY720 treatment has been shown to alter the biology of multiple cell types beyond conventional T cells, including increasing endothelial permeability (Shea et al., 2010; Oo et al., 2011), potentially altering inflammatory cell recruitment, as well as potently inhibiting Treg function (Wolf et al., 2009). Likewise, anti-Thy1 antibody likely targets multiple cell types, including Tregs. Parabiosis allowed us to define the function of Th2 Trm and circulating memory Th2 cells in an unmanipulated re-challenge response, demonstrating that both Th2 cell subsets contribute to the recall response and perform distinct functions. While our parabiosis model allows us to define the role of memory Th2 cell subsets *in vivo*, one limitation of our study is that we have not defined the additional lung-resident changes in HDM-memory mice that promote the recall response to HDM. Specifically, the niches maintaining Th2 Trm within the lung remain poorly delineated. Notably, there appear to be differences in the tissue environment supporting distinct types of Trm. Lung CD8^+^ Trm have been shown to persist in “repair-associated memory depots” independently of conventional inducible bronchus-associated lymphoid tissue (iBALT) (Takamura et al., 2016). Similarly, Th1 Trm in the female reproductive tract have also been shown to persist in “macrophage lymphocyte clusters” independently of tertiary lymphoid structures (Iijima and Iwasaki, 2014). In contrast, memory Th2 cells persisting in the lung of mice after allergic inflammation have been shown to localize to iBALT-like structures, supported by IL-7 from lymphatic endothelial cells (Shinoda et al., 2016). Future studies will be needed to define the unique features of the peribronchial niche supporting the development, maintenance, and function of Th2 Trm *in vivo*.

In summary, we have used a HDM model of allergic airway disease, parabiosis, and transcriptional analysis to define the function and transcriptional profiles of tissue-resident and circulating memory Th2 cells. We demonstrate that Th2 Trm within the lung and circulating memory Th2 cells are transcriptionally distinct subsets that exhibit non-overlapping trafficking patterns during homeostasis. Upon allergen re-challenge, Th2 Trm and circulating memory Th2 cells both contribute to the increased numbers of Th2 cells and type 2 cytokine expression in the lung and perform non-redundant functions *in vivo* (Fig. 5F). Consequently, Th2 Trm represent a unique Th2 cell subset with a distinct tissue adaptive gene signature that are required for the full manifestation of allergic airway disease *in vivo*. Defining the unique mechanisms supporting the development and maintenance of Th2 Trm will be critical to developing novel therapeutic approaches for allergic asthma.

## Methods

### Mice

C57BL/6J (Wild type) and CD45.1 CRL mice were obtained from Charles River Laboratories. *Rag2*-deficient and C57BL6/J control mice were obtained from Jackson laboratories. FoxP3*^YFPCre^* mice were obtained from Jackson laboratories and are bred and maintained within our facility. All mice analyzed were sex and aged matched (6-12 weeks old). All mice were bred and maintained in specific-pathogen-free conditions at the animal facility of the Massachusetts General Hospital and were used under a study protocol approved by Massachusetts General Hospital Subcommittee on Research Animal Care.

### Mouse treatments

To induce allergic airway inflammation, mice were anesthetized with intramuscular injection of Ketamine/Xylazine (Patterson Veterinary) and sensitized via intranasal administration (i.n.) with 10 μg HDM (*Dermatophagoides pteronyssinus* extracts, Greer Laboratories) in 40 μl of sterile PBS on day 0. On day 7, mice were challenged with 10 μg of HDM via i.n. route daily for five days. 6-12 weeks later mice were either left untreated or re-challenged with 10 μg of HDM via i.n. route once or 10 μg of *Alternaria Alternata* extract (Greer Laboratories). For Th2 proliferation studies, 1-2 mg of BrdU (Thermo Fisher Scientific) was injected intraperitoneally and tissue was harvested 2 hours later.

### Parabiosis surgery

CD45.2 C57BL/6J mice were sensitized and challenged with HDM as outlined and rested for 3-4 weeks to generate memory Th2 cells in vivo. CD45.2 HDM-memory C57BL/6J mice and naïve CD45.1 B6 mice were surgically conjoined to achieve parabiosis. Specifically, mice underwent hair removal along opposite lateral flanks with the use of hair clippers and depilatory cream. Skin was then wiped clean of fur and sterilized with topical betadine solution. Mirrored incisions were then made on the lateral aspects of both mice from forelimb to hindlimb. 4.0 sutures were placed around the olecranon and knee joints to secure the upper and lower extremities of the mice. Dorsal and ventral skin was approximated with a running 4.0 suture and surgical staples. At the end of the surgery, mice received subcutaneous enrofloxacin antibiotic as well as buprenorphine and flunixin for pain control. Enrofloxacin antibiotic was subsequently administered via drinking water for 2 weeks. Subcutaneous buprenorphine and flunixin was administered as needed every 12 hours for 48 hours. Parabionts were both challenged with 10 μg of HDM i.n. BAL, lung, and mediastinal lymph nodes were collected at 72 hours.

### Tissue harvest and leukocyte preparation

Prior to tissue harvest, mice were anesthetized with Ketamine/Xylazine (Patterson Veterinary) and injected intravenously with 3 μg of fluorophore-labelled anti-CD45 (30-F11, Biolegend) monoclonal antibody through the retro-orbital sinus to label intravascular leukocytes. Three minutes after anti-CD45 antibody injection, the trachea was exposed, cannulated and bronchoalveolar lavage was performed by infusing 3 ml of cold PBS with 0.5 mM EDTA. Lung lobes and mLN were subsequently removed, minced with scissors and digested at 37°C for 20-30 minutes in digestion buffer (0.52 U/ml Liberase TM [Roche] and 60 U/ml DNase I [Roche] in RPMI 1640 [Cellgro] with 5% fetal bovine serum). Digested tissue was strained through a 70-μm filter to generate a single cell suspension.

### Flow cytometry

Single cells were incubated with anti-mouse CD16/32 (93, TruStain fcX, BioLegend) to block Fc receptors. Staining was performed with Fixable Viability Dye eF780 or eF506 (eBioscience), to identify dead cells, and the following fluorochrome-conjugated anti-mouse monoclonal antibodies (mAbs): CD3e-AF488 (145-2C11), CD4-PeCy7 (GK1.5), Foxp3-APC (FJK-16s), GATA3-PE (L50-823), CD45-APC-eFluor780 (30-F11), CD11c-BV605 (N418), Siglec-F-PE (E50-2440), CD11b-BUV737 (M1/70), MHCII-PeCy7 (M5/114.15.2), CD45.2-BUV395 (104), CD62L-FITC (MEL14), BrdU-AF647 (3D4), ST2-BV421 (U29-93), CD127-PeCy7 (A7R34), Thy1.2-PerCP-eFluor710 (30-H12). Flow cytometric analysis was performed using a LSR Fortessa X-20 flow cytometer (BD Biosciences) and FlowJo software (Tree Star). Intracellular staining was performed using eBioscience Fixation Permeabilization buffers. BrdU staining was performed following the BDBiosciences manufacture’s protocol.

### *Ex vivo* T cell re-stimulation

Mediastinal lymph nodes were harvested and single cell suspensions obtained. 0.25 x 10^6^ total lymph node cells were cultured in 250 ul of RPMI 1640 media (Cellgro) supplemented with 10% (v/v) heat-inactivated fetal bovine serum (FBS, Sigma), 50 μM 2-mercaptoethanol, 2 mM Glutamax (Gibco), 5 mM HEPES, 1 mM sodium pyruvate, 0.1 mM non-essential amino acids, 100 U/mL penicillin-streptomycin (all from Lonza) and stimulated with 25 μg/ml HDM (Greer Laboratories) for 72 h at 37°C, 5% CO2. Supernatant culture cytokine levels for IL-5 (Biolegend) and IL-13 (eBioscience) were measured by enzyme-linked immunosorbent assay (ELISA) utilizing Biolegend and eBioscience kits, respectively, and according to the manufacturer’s instructions.

### qPCR RNA levels

RNA was extracted from lung tissue in Trizol using RNeasy Mini kit (Qiagen) and cDNA was reverse-transcribed using SuperScript III First Strand (Invitrogen) following manufacturer’s guidelines. qPCR reactions were performed on a LightCylcer 96 Instrument (Roche) using FastStart Essential DNA Green Master (Roche) and normalized to *B2m* or *Gapdh* using the following primers: *Il5*, 5’-CTC TGT TGA CAA GCA ATG AGA CG-3’ (forward) and 5’-TCT TCA GTA TGT CTA GCC CCT G-3’ (reverse); *Il13*, 5’-CCT GGC TCT TGC TTG CCTT-3’ (forward) and 5’-GGT CTT GTG TGA TGT TGC TCA-3’ (reverse); *Ccl11*, 5’-TCC ACA GCG CTT CTA TTC CTG -3’ (forward) and 5’-GGA GCC TGG GTG AGC CA -3’ (reverse); *Ccl24*, 5’-ATT CTG TGA CCA TCC CCT CAT -3’ (forward) and 5’-TGT ATG TGC CTC TGA ACC CAC -3’ (reverse); *Ccl17*, 5’-CAG GGA TGC CAT CGT GTT TC -3’ (forward) and 5’-CAC CAA TCT GAT GGC CTT CTT -3’ (reverse); *Ccl22*, 5’-TAC ATC CGT CAC CCT CTG CC -3’ (forward) and 5’-CGG TTA TCA AAA CAA CGC CAG -3’ (reverse); *B2m*, 5’-CCC GTT CTT CAG CAT TTG GA -3’ (forward) and 5’-CCG AAC ATA CTG AAC TGC TAC GTA A -3’ (reverse); *Gapdh*, 5’-GGC AAA TTC AAC GGC ACA GT-3’ (forward) and 5’-AGA TGG TGA TGG GCT TCC C-3’ (reverse). Results indicated as relative expression indicate copies per *B2m* or *Gapdh*.

### Histology

Lung samples were fixed in buffered 10% formalin solution. Paraffin-embedded sections were cut (5 mm) and stained with hematoxylin and eosin (H&E) or periodic acid-Schiff (PAS). Lung H&E histology scoring was performed by a Pathologist from the Harvard Pathology Core who was blinded to the groups. Goblet cells were enumerated from PAS-stained sections using a numerical scoring system (0, <5% goblet cells; 1, 5-25%; 2, 25-50%; 3, 50-75%; 4, >75%). 20-50 airways per mouse were evaluated and the sum of goblet cell scoring from each mouse was divided by the number of airways and presented as a mucus score as previously described (Patel et al., 2019). All images are shown at 10x magnification.

### Measurement of airway resistance

Airway resistance was measured using the SCIREQ flexiVent system. Mice were anesthetized with Ketamine/Xylazine and intubated via tracheotomy. The mice were placed on a ventilator and paralyzed with pancuonium (1 mg/kg). After 5 minutes, the mice were first challenged with PBS and then progressively challenged with methacholine (12.5 mg/mL, 25 mg/ml and 50 mg/mL) and airway resistance was measured using the flexiVent software system.

### RNA-seq analysis

Sequencing was performed on Illumina HiSeq 2500 instrument, resulting in approximately 30 million of 50 bp reads per sample. Sequencing reads were mapped in a splice-aware fashion to the mouse reference transcriptome (mm9 assembly) using STAR (Dobin et al., 2013). Read counts over transcripts were calculated using HTSeq based on the Ensembl annotation for GRCm37/mm9 assembly (Anders et al., 2015). For differential expression analysis we used the EdgeR method and classified genes as differentially expressed based on the cutoffs of 2-fold change in expression value and false discovery rates (FDR) below 0.05 (Robinson et al., 2009).

### Statistical analysis

All statistical analyses were performed using GraphPad Prism 7 software. Statistical parameters are reported in the Figure Legends but generally *P* values were determined for multiple comparisons by one-way ANOVA analysis with Holm-Sidak’s testing. Unpaired or paired two-tailed Student’s *t* test was used for two group comparisons for normally distributed data. Mann-Whitney U testing was used for two group comparisons for non-normally distributed data. In all Figures, statistical significance is indicated (\**P*<0.05; \*\**P*<0.01; \*\*\**P*<0.001; \*\*\*\**P*<0.0001). No statistical method was used to predetermine sample size. The number of mice used in each experiment to reach statistical significance was determined on the basis of preliminary data. No animals were excluded from the analyses. Blinding was used for lung H&E history scoring. Data met assumptions of statistical methods used and variance was similar between groups that were statistically compared.

## Acknowledgements

This work was supported by grants from the National Institutes of Health R01 AI040618 (A.D.L) and the Training Program in Pulmonary Immunology and Allergy T32 HL116275 (A.D.L.) as well as K08 HL140173 (R.A.R) and the National Institutes of Health P30 DK040561 (R.I.S.).

Cytometric findings reported here were performed in the MGH Department of Pathology Flow and Image Cytometry Research Core, which obtained support from the NIH Shared Instrumentation program with grants 1S10OD012027-01A1, 1S10OD016372-01, 1S10RR020936-01, and 1S10RR023440-01A1.

We thank Dr. Hui Zheng from the MGH Biostatistics Center for thoughtful input on statistical analyses. We thank members of the MGH Next Generation Sequencing Core facility and Flow Cytometry Core facility for technical assistance.

## Author contribution

R.A.R designed, performed, analyzed and interpreted experiments and wrote the manuscript. K.N. performed experiments, collected and assembled data. M.C. and R.I.S assisted with RNA-seq data analysis and presentation as well as manuscript preparation. A.D.L. conceptualized and designed the study, analyzed and interpreted data, and contributed to manuscript preparation.

---

**Supplemental Figure 1.**
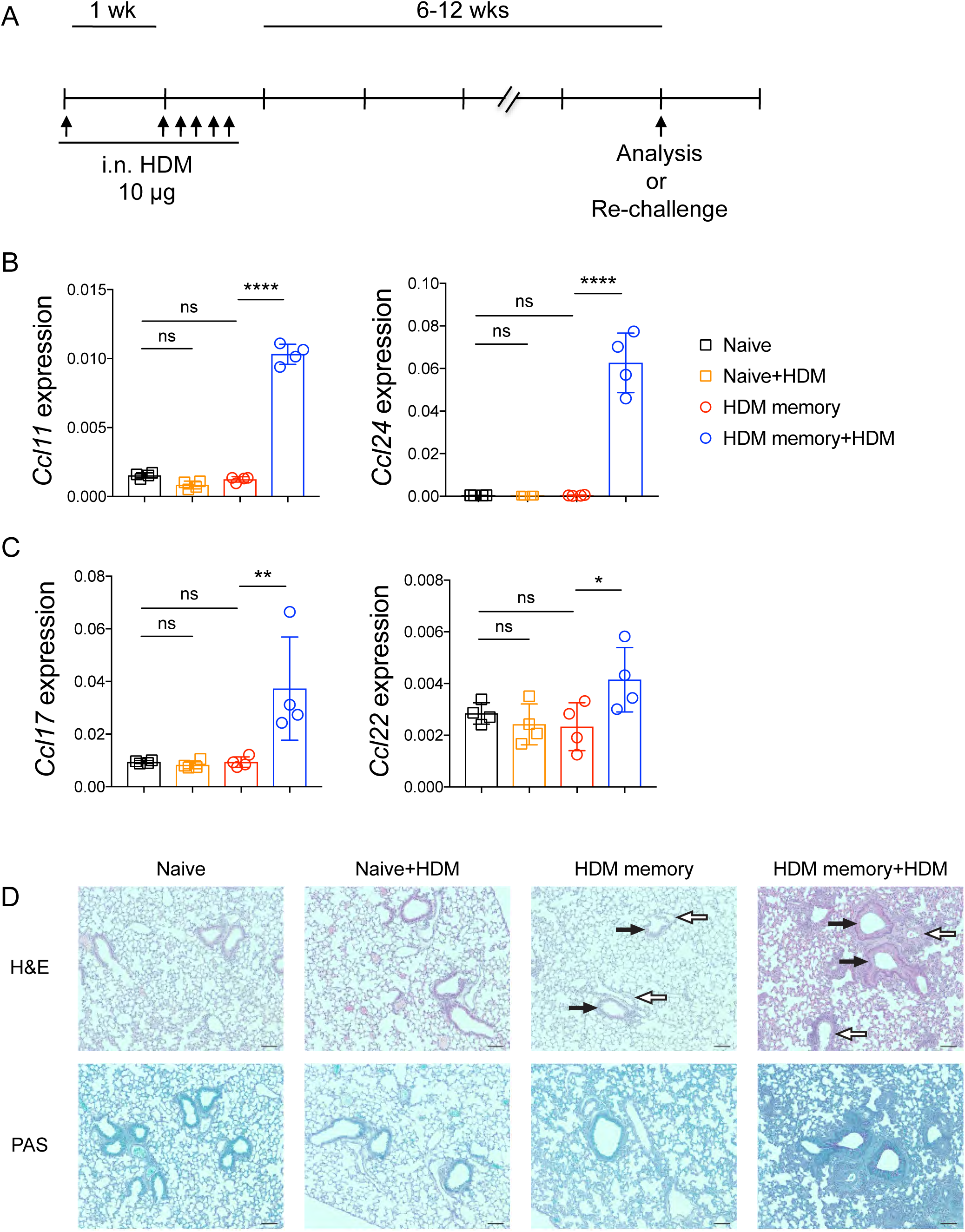
**Allergen re-challenge promotes type 2 chemokine expression, lung inflammation, and mucus metaplasia.** C57BL/6 mice were sensitized with 10 μg of intranasal (i.n.) HDM followed by 10 μg daily challenges of i.n. HDM on days 7-11. After 6-12 weeks of rest, HDM-memory mice were left untreated or re-challenged with a single dose of i.n. HDM followed by tissue harvest 72 hours later. (A) Schematic of i.n. HDM treatment. (B) Lung *Ccl11* and *Ccl24* relative RNA levels assessed via qPCR. (C) Lung *Ccl17* and *Ccl22* relative RNA levels assessed via qPCR. (D) Representative images of lung H&E and PAS staining in indicated groups. White arrows indicate blood vessels, black arrows indicate airways. Scale bars represent 100 μm. Representative data show individual mice with mean ± SEM from one of two independent experiments with 3-4 mice per group. For statistical analysis, one-way ANOVA analysis with Holm-Sidak’s testing was used for statistical analysis of multiple groups. *P < 0.05; **P < 0.01; ***P < 0.001.

**Supplemental Figure 2.**
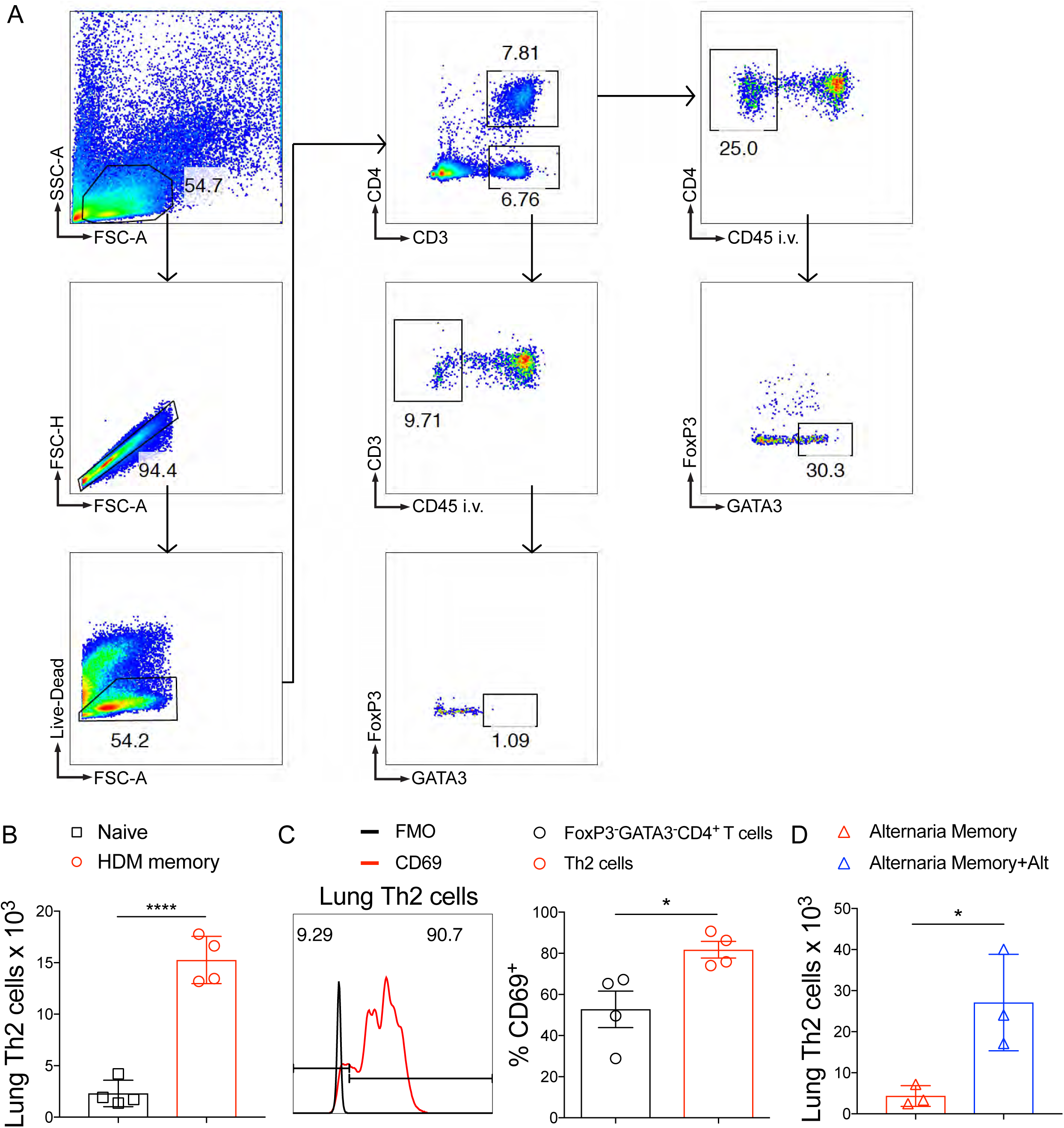
**Identification of tissue-resident memory Th2 cells via flow cytometry.** C57BL/6 mice were sensitized and challenged with i.n. HDM then rested for 6-12 weeks followed by lung tissue harvest. (A) Representative flow cytometry on lung cells in HDM-memory mice to identify Th2 cells (anti-CD45 i.v. unlabeled, FoxP3^-^ GATA3^+^CD4^+^ T cells). (B) Quantitation of lung Th2 cells (anti-CD45 i.v. unlabeled, FoxP3^-^GATA3^+^CD4^+^ T cells) in naive and HDM-memory mice via flow cytometry. (C) Representative histogram demonstrating CD69 staining (and fluorescence minus one control staining) of lung Th2 cells from HDM-memory mice (left panel) with quantitation of percent CD69^+^ cells from lung Th2 cells and FoxP3^-^GATA3^-^CD4^+^ T cell populations (right panel). (D) C57BL/6 mice were sensitized and challenged with i.n. *Alternaria*, rested for 6-8 weeks, and left untreated or re-challenged with a single dose of i.n. *Alternaria*. Quantitation of lung Th2 cells (anti-CD45 i.v. unlabeled, FoxP3^-^GATA3^+^CD4^+^ T cells) in indicated groups. Representative data show individual mice with mean ± SEM from one of two independent experiments with 3-4 mice per group. For statistical analysis, one-way ANOVA analysis with Holm-Sidak’s testing was used for statistical analysis of multiple groups. For comparison of two groups, a two-tailed t test was performed. *P < 0.05; **P < 0.01; ***P < 0.001.

**Supplemental Figure 3.**
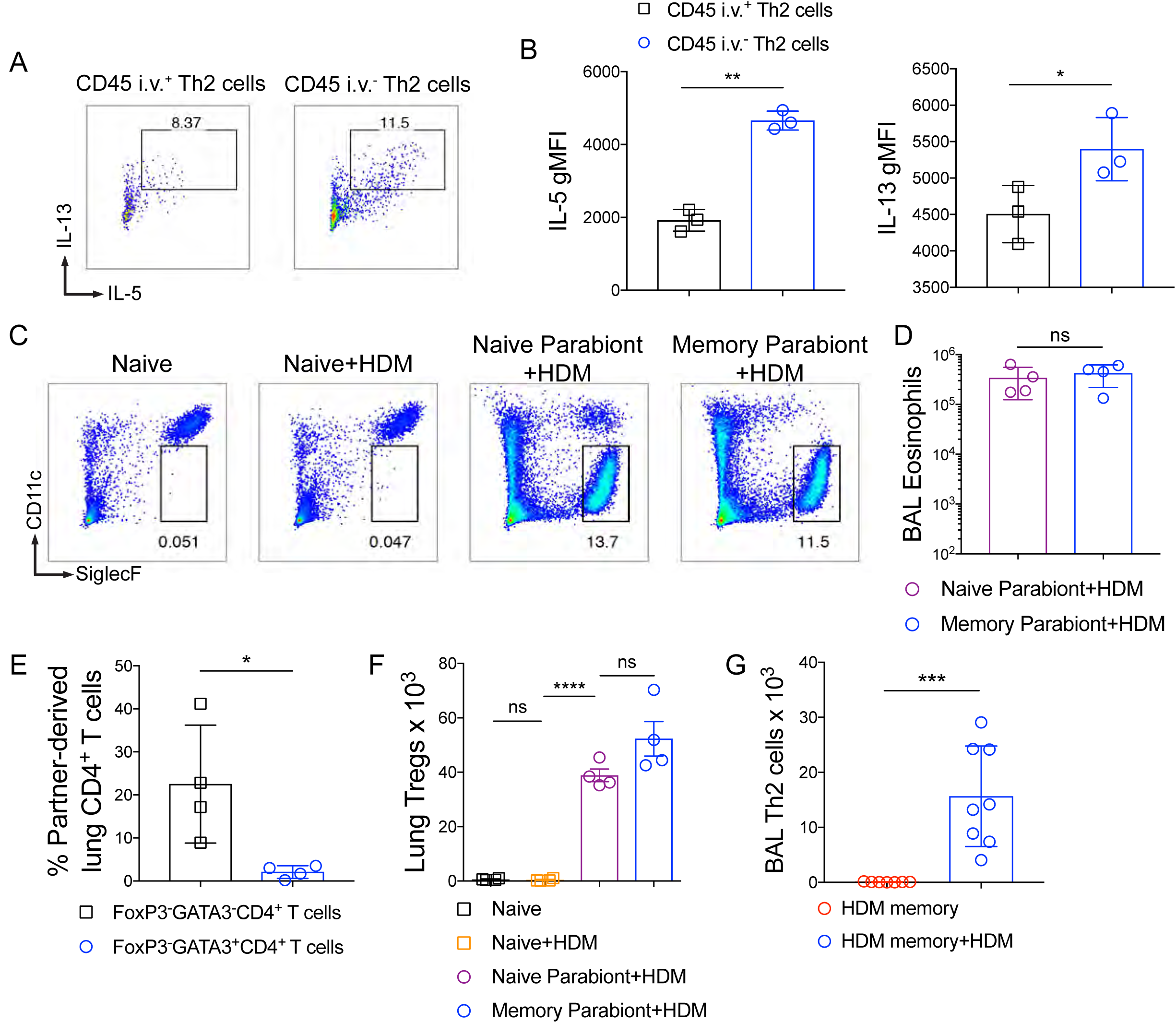
**Function and trafficking of tissue-resident and circulating memory Th2 cells in HDM-memory mice.** (A-B) C57BL/6 mice were sensitized and challenged with i.n. HDM and rested for 6-12 weeks. (A) Lung cells from HDM-memory mice underwent CD4^+^ T cell negative selection followed by treatment with anti-CD3 and anti-CD28 for 12-16 hours. Representative flow cytometry of anti-CD45 i.v. unlabeled (intraparenchymal) and labeled (intravascular) Th2 cells after intracellular cytokine staining for IL-5 and IL-13. (B) Geometric mean fluorescence intensity (gMFI) for IL-5 and IL-13 expression in indicated groups. (C-D) CD45.2 HDM-memory mice were surgically conjoined to CD45.1 naïve mice. After 3-4 weeks both parabionts received a single dose of i.n. HDM with harvest of mLN and lung after 72 hours. (C) Representative flow cytometry of lung eosinophils (anti-CD45 i.v. unlabeled, CD11c^lo^Siglec-F^+^ cells) from indicated groups. (D) Quantitation of BAL eosinophils from indicated groups. (E) CD45.2 and CD45.1 HDM-memory mice were surgically conjoined. After 3-4 weeks lungs from both parabionts were harvested. Quantitation of percent partner-derived lung memory CD4^+^ T cells (anti-CD45 i.v. unlabeled, FoxP3^-^GATA3^-^CD4^+^ T cells and FoxP3^-^ GATA3^+^CD4^+^ T cells) assessed via flow cytometry. (F) CD45.2 HDM-memory mice were surgically conjoined to CD45.1 naïve mice. After 3-4 weeks both parabionts received a single dose of i.n. HDM with harvest of lung after 72 hours. Lung FoxP3^+^CD4^+^ T cells (Tregs) were quantitated via flow cytometry. (G) Individual HDM-memory mice were left untreated or received a single dose of i.n. HDM with BAL after 72 hours. BAL Th2 cells were quantitated via flow cytometry. Representative data show individual mice with mean ± SEM from one of two independent experiments with 3-4 mice per group (A-B), or one of three independent experiments with 3-4 mice per group (C-D, F), or cumulative data from two independent experiments (E, G). For statistical analysis, a two-tailed t test was performed for parametric data (A-B, G). One-way ANOVA analysis with Holm-Sidak’s testing was used for statistical analysis of multiple groups with paired two-tailed t tests for comparison on naive and memory parabiont groups (C-D, F). A two-tailed Mann-Whitney U test was performed for non-parametric data (E). *P < 0.05; **P < 0.01.

## References

1. Afshar, R., J.P. Strassner, E. Seung, B. Causton, J.L. Cho, R.S. Harris, D.L. Hamilos, B.D. Medoff, and A.D. Luster. 2013. Compartmentalized chemokine-dependent regulatory T-cell inhibition of allergic pulmonary inflammation. J. Allergy Clin. Immunol. 131:1644–1652.e4. doi:10.1016/j.jaci.2013.03.002.

2. Anders, S., P.T. Pyl, and W. Huber. 2015. HTSeq-A Python framework to work with high-throughput sequencing data. Bioinformatics. 31:166–169. doi:10.1093/bioinformatics/btu638.

3. Anderson, K.G., K. Mayer-Barber, H. Sung, L. Beura, B.R. James, J.J. Taylor, L. Qunaj, T.S. Griffith, V. Vezys, D.L. Barber, and D. Masopust. 2014. Intravascular staining for discrimination of vascular and tissue leukocytes. Nat. Protoc. 9:209–222. doi:10.1038/nprot.2014.005.

4. Angela, M., Y. Endo, H.K. Asou, T. Yamamoto, D.J. Tumes, H. Tokuyama, K. Yokote, and T. Nakayama. 2016. Fatty acid metabolic reprogramming via mTOR-mediated inductions of PPARγ directs early activation of T cells. Nat. Commun. 7:1–15. doi:10.1038/ncomms13683.

5. Arbes, S.J., P.J. Gergen, B. Vaughn, and D.C. Zeldin. 2007. Asthma cases attributable to atopy: Results from the Third National Health and Nutrition Examination Survey. J. Allergy Clin. Immunol. 120:1139–1145. doi:10.1016/j.jaci.2007.07.056.

6. Beura, L.K., N.J. Fares-Frederickson, E.M. Steinert, M.C. Scott, E.A. Thompson, K.A. Fraser, J.M. Schenkel, V. Vezys, and D. Masopust. 2019. CD4+ resident memory T cells dominate immunosurveillance and orchestrate local recall responses. J. Exp. Med. 216:1214–1229. doi:10.1084/jem.20181365.

7. Beura, L.K., J.S. Mitchell, E.A. Thompson, J.M. Schenkel, J. Mohammed, S. Wijeyesinghe, R. Fonseca, B.J. Burbach, H.D. Hickman, V. Vezys, B.T. Fife, and D. Masopust. 2018. Intravital mucosal imaging of CD8 + resident memory T cells shows tissue-autonomous recall responses that amplify secondary memory article. Nat. Immunol. 19:173–182. doi:10.1038/s41590-017-0029-3.

8. Bromley, S.K., S. Yan, M. Tomura, O. Kanagawa, and A.D. Luster. 2013. Recirculating Memory T Cells Are a Unique Subset of CD4 + T Cells with a Distinct Phenotype and Migratory Pattern . J. Immunol. 190:970–976. doi:10.4049/jimmunol.1202805.

9. Carbone, F.R. 2015. Tissue-Resident Memory T Cells and Fixed Immune Surveillance in Nonlymphoid Organs. J. Immunol. 195:17–22. doi:10.4049/jimmunol.1500515.

10. Causton, B., A. Pardo-Saganta, J. Gillis, K. Discipio, T. Kooistra, J. Rajagopal, R.J. Xavier, J.L. Cho, and B.D. Medoff. 2018. CARMA3 mediates allergic lung inflammation in response to alternaria alternata. Am. J. Respir. Cell Mol. Biol. 59:684–694. doi:10.1165/rcmb.2017-0181OC.

11. Chen, T., C.A. Tibbitt, X. Feng, J.M. Stark, L. Rohrbeck, L. Rausch, S.K. Sedimbi, M.C.I. Karlsson, B.N. Lambrecht, G.B. Karlsson Hedestam, R.W. Hendriks, B.J. Chambers, S. Nylén, and J.M. Coquet. 2017. PPAR-promotes type 2 immune responses in allergy and nematode infection. Sci. Immunol. 2:1–11. doi:10.1126/sciimmunol.aal5196.

12. Dobin, A., C.A. Davis, F. Schlesinger, J. Drenkow, C. Zaleski, S. Jha, P. Batut, M. Chaisson, and T.R. Gingeras. 2013. STAR: Ultrafast universal RNA-seq aligner. Bioinformatics. 29:15–21. doi:10.1093/bioinformatics/bts635.

13. Endo, Y., K. Hirahara, T. Iinuma, K. Shinoda, D.J. Tumes, H.K. Asou, N. Matsugae, K. Obata-Ninomiya, H. Yamamoto, S. Motohashi, K. Oboki, S. Nakae, H. Saito, Y. Okamoto, and T. Nakayama. 2015. The Interleukin-33-p38 kinase axis confers memory T helper 2 cell pathogenicity in the airway. Immunity. 42:294–308. doi:10.1016/j.immuni.2015.01.016.

14. Endo, Y., C. Iwamura, M. Kuwahara, A. Suzuki, K. Sugaya, D.J. Tumes, K. Tokoyoda, H. Hosokawa, M. Yamashita, and T. Nakayama. 2011. Eomesodermin controls interleukin-5 production in memory T helper 2 cells through inhibition of activity of the transcription factor GATA3. Immunity. 35:733–745. doi:10.1016/j.immuni.2011.08.017.

15. Evans, C.M., D.S. Raclawska, F. Ttofali, D.R. Liptzin, A.A. Fletcher, D.N. Harper, M.A. McGing, M.M. McElwee, O.W. Williams, E. Sanchez, M.G. Roy, K.N. Kindrachuk, T.A. Wynn, H.K. Eltzschig, M.R. Blackburn, M.J. Tuvim, W.J. Janssen, D.A. Schwartz, and B.F. Dickey. 2015. The polymeric mucin Muc5ac is required for allergic airway hyperreactivity. Nat. Commun. 6:1–11. doi:10.1038/ncomms7281.

16. Faustino, L., D.M. da Fonseca, M.C. Takenaka, L. Mirotti, E.B. Florsheim, M.G. Guereschi, J.S. Silva, A.S. Basso, and M. Russo. 2013. Regulatory T Cells Migrate to Airways via CCR4 and Attenuate the Severity of Airway Allergic Inflammation. J. Immunol. 190:2614–2621. doi:10.4049/jimmunol.1202354.

17. Galkina, E., J. Thatte, V. Dabak, M.B. Williams, K. Ley, and T.J. Braciale. 2005. Preferential migration of effector CD8+ T cells into the interstitium of the normal lung. J. Clin. Invest. 115:3473–3483. doi:10.1172/JCI24482.

18. Gerlach, C., E.A. Moseman, S.M. Loughhead, D. Alvarez, A.J. Zwijnenburg, L. Waanders, R. Garg, J.C. de la Torre, and U.H. von Andrian. 2016. The Chemokine Receptor CX3CR1 Defines Three Antigen-Experienced CD8 T Cell Subsets with Distinct Roles in Immune Surveillance and Homeostasis. Immunity. 45:1270–1284. doi:10.1016/j.immuni.2016.10.018.

19. Glennie, N.D., V.A. Yeramilli, D.P. Beiting, S.W. Volk, C.T. Weaver, and P. Scott. 2015. Skin-resident memory CD4+ T cells enhance protection against Leishmania major infection. J. Exp. Med. 212:1405–1414. doi:10.1084/jem.20142101.

20. Griffith, J.W., C.L. Sokol, and A.D. Luster. 2014. Chemokines and Chemokine Receptors: Positioning Cells for Host Defense and Immunity. Annu. Rev. Immunol. 32:659–702. doi:10.1146/annurev-immunol-032713-120145.

21. Guo, L., Y. Huang, X. Chen, J. Hu-Li, J.F. Urban, and W.E. Paul. 2015. Innate immunological function of T H2 cells in vivo. Nat. Immunol. 16:1051–1059. doi:10.1038/ni.3244.

22. Guo, L., G. Wei, J. Zhu, W. Liao, W.J. Leonard, K. Zhao, and W. Paul. 2009. IL-1 family members and STAT activators induce cytokine production by Th2, Th17, and Th1 cells. Proc. Natl. Acad. Sci. U. S. A. 106:13463–13468. doi:10.1073/pnas.0906988106.

23. Hofmann, M., V. Brinkmann, and H.G. Zerwes. 2006. FTY720 preferentially depletes naive T cells from peripheral and lymphoid organs. Int. Immunopharmacol. 6:1902– 1910. doi:10.1016/j.intimp.2006.07.030.

24. Hombrink, P., C. Helbig, R.A. Backer, B. Piet, A.E. Oja, R. Stark, G. Brasser, A. Jongejan, R.E. Jonkers, B. Nota, O. Basak, H.C. Clevers, P.D. Moerland, D. Amsen, and R.A.W. Van Lier. 2016. Programs for the persistence, vigilance and control of human CD8 + lung-resident memory T cells. Nat. Immunol. 17:1467– 1478. doi:10.1038/ni.3589.

25. Hondowicz, B.D., D. An, J.M. Schenkel, K.S. Kim, H.R. Steach, A.T. Krishnamurty, G.J. Keitany, E.N. Garza, K.A. Fraser, J.J. Moon, W.A. Altemeier, D. Masopust, and M. Pepper. 2016. Interleukin-2-Dependent Allergen-Specific Tissue-Resident Memory Cells Drive Asthma. Immunity. 44:155–166. doi:10.1016/j.immuni.2015.11.004.

26. Huang, D.W., B.T. Sherman, and R.A. Lempicki. 2009. Systematic and integrative analysis of large gene lists using DAVID bioinformatics resources. Nat. Protoc. 4:44–57. doi:10.1038/nprot.2008.211.

27. Ichikawa, T., K. Hirahara, K. Kokubo, M. Kiuchi, A. Aoki, Y. Morimoto, J. Kumagai, A. Onodera, N. Mato, D.J. Tumes, Y. Goto, K. Hagiwara, Y. Inagaki, T. Sparwasser, K. Tobe, and T. Nakayama. 2019. CD103hi Treg cells constrain lung fibrosis induced by CD103lo tissue-resident pathogenic CD4 T cells. Nat. Immunol. 20:1469–1480. doi:10.1038/s41590-019-0494-y.

28. Iijima, N., and A. Iwasaki. 2014. A local macrophage chemokine network sustains protective tissue-resident memory CD4 T cells. Science (80-.). 346:93–98. doi:10.1126/science.1257530.

29. Jameson, S.C., and D. Masopust. 2018. Understanding Subset Diversity in T Cell Memory. Immunity. 48:214–226. doi:10.1016/j.immuni.2018.02.010.

30. Johansson, M.W. 2014. Activation states of blood eosinophils in asthma. Clin. Exp. Allergy. 44:482–498. doi:10.1111/cea.12292.

31. Lambrecht, B.N., and H. Hammad. 2015. The immunology of asthma. Nat. Immunol. 16:45–56. doi:10.1038/ni.3049.

32. Lambrecht, B.N., H. Hammad, and J. V. Fahy. 2019. The Cytokines of Asthma. Immunity. 50:975–991. doi:10.1016/j.immuni.2019.03.018.

33. Li, B.W.S., M.J.W. de Bruijn, I. Tindemans, M. Lukkes, A. KleinJan, H.C. Hoogsteden, and R.W. Hendriks. 2016. T cells are necessary for ILC2 activation in house dust mite-induced allergic airway inflammation in mice. Eur. J. Immunol. 46:1392–1403. doi:10.1002/eji.201546119.

34. Lukacs, N.W. 2001. Role of chemokines in the pathogenesis of asthma. Nat. Rev. Immunol. 1:108–116. doi:10.1038/35100503.

35. Mackay, L.K., M. Minnich, N.A.M. Kragten, Y. Liao, B. Nota, C. Seillet, A. Zaid, K. Man, S. Preston, D. Freestone, A. Braun, E. Wynne-Jones, F.M. Behr, R. Stark, D.G. Pellicci, D.I. Godfrey, G.T. Belz, M. Pellegrini, T. Gebhardt, M. Busslinger, W. Shi, F.R. Carbone, R.A.W. Van Lier, A. Kallies, and K.P.J.M. Van Gisbergen. 2016. Hobit and Blimp1 instruct a universal transcriptional program of tissue residency in lymphocytes. Science (80-.). 352:459–463. doi:10.1126/science.aad2035.

36. MacKay, L.K., A. Rahimpour, J.Z. Ma, N. Collins, A.T. Stock, M.L. Hafon, J. Vega-Ramos, P. Lauzurica, S.N. Mueller, T. Stefanovic, D.C. Tscharke, W.R. Heath, M. Inouye, F.R. Carbone, and T. Gebhardt. 2013. The developmental pathway for CD103+ CD8+ tissue-resident memory T cells of skin. Nat. Immunol. 14:1294– 1301. doi:10.1038/ni.2744.

37. Martinez-Gonzalez, I., L. Mathä, C.A. Steer, M. Ghaedi, G.F.T. Poon, and F. Takei. 2016. Allergen-Experienced Group 2 Innate Lymphoid Cells Acquire Memory-like Properties and Enhance Allergic Lung Inflammation. Immunity. 45:198–208. doi:10.1016/j.immuni.2016.06.017.

38. Martinez-Gonzalez, I., L. Mathä, C.A. Steer, and F. Takei. 2017. Immunological Memory of Group 2 Innate Lymphoid Cells. Trends Immunol. 38:423–431. doi:10.1016/j.it.2017.03.005.

39. McKnight, C.G., J.A. Jude, Z. Zhu, R.A. Panettieri, and F.D. Finkelman. 2017. House dust mite-induced allergic airway disease is independent of IgE and FceRIa. Am. J. Respir. Cell Mol. Biol. 57:674–682. doi:10.1165/rcmb.2016-0356OC.

40. Mikhak, Z., M. Fukui, A. Farsidjani, B.D. Medoff, A.M. Tager, and A.D. Luster. 2009. Contribution of CCR4 and CCR8 to antigen-specific TH2 cell trafficking in allergic pulmonary inflammation. J. Allergy Clin. Immunol. 123:67–73.e3. doi:10.1016/j.jaci.2008.09.049.

41. Mikhak, Z., J.P. Strassner, and A.D. Luster. 2013. Lung dendritic cells imprint T cell lung homing and promote lung immunity through the chemokine receptor CCR4. J. Exp. Med. 210:1855–1869. doi:10.1084/jem.20130091.

42. Milner, J.J., C. Toma, B. Yu, K. Zhang, K. Omilusik, A.T. Phan, D. Wang, A.J. Getzler, T. Nguyen, S. Crotty, W. Wang, M.E. Pipkin, and A.W. Goldrath. 2017. Runx3 programs CD8+ T cell residency in non-lymphoid tissues and tumours. Nature. 552:253–257. doi:10.1038/nature24993.

43. Minutti, C.M., S. Drube, N. Blair, C. Schwartz, J.C. McCrae, A.N. McKenzie, T. Kamradt, M. Mokry, P.J. Coffer, M. Sibilia, A.J. Sijts, P.G. Fallon, R.M. Maizels, and D.M. Zaiss. 2017. Epidermal Growth Factor Receptor Expression Licenses Type-2 Helper T Cells to Function in a T Cell Receptor-Independent Fashion. Immunity. 47:710–722.e6. doi:10.1016/j.immuni.2017.09.013.

44. Nobs, S.P., S. Natali, L. Pohlmeier, K. Okreglicka, C. Schneider, M. Kurrer, F. Sallusto, and M. Kopf. 2017. PPARγ in dendritic cells and T cells drives pathogenic type-2 effector responses in lung inflammation. J. Exp. Med. 214:3015–3035. doi:10.1084/jem.20162069.

45. Oja, A.E., B. Piet, C. Helbig, R. Stark, D. Van Der Zwan, H. Blaauwgeers, E.B.M. Remmerswaal, D. Amsen, R.E. Jonkers, P.D. Moerland, M.A. Nolte, R.A.W. Van Lier, and P. Hombrink. 2018. Trigger-happy resident memory CD4 + T cells inhabit the human lungs. Mucosal Immunol. 11:654–667. doi:10.1038/mi.2017.94.

46. Onodera, A., K. Kokubo, and T. Nakayama. 2018. Epigenetic and Transcriptional Regulation in the Induction, Maintenance, Heterogeneity, and Recall-Response of Effector and Memory Th2 Cells. Front. Immunol. 9:2929. doi:10.3389/fimmu.2018.02929.

47. Oo, M.L., D.K. Han, T. Hla, M.L. Oo, S. Chang, S. Thangada, M. Wu, and K. Rezaul. 2011. Engagement of S1P 1-degradative mechanisms leads to vascular leak in mice Find the latest version : Engagement of S1P 1 -degradative mechanisms leads to vascular leak in mice. J. Clin. Invest. 121:2290–2300. doi:10.1172/JCI45403.2290.

48. Pan, Y., T. Tian, C.O. Park, S.Y. Lofftus, S. Mei, X. Liu, C. Luo, J.T. O’Malley, A. Gehad, J.E. Teague, S.J. Divito, R. Fuhlbrigge, P. Puigserver, J.G. Krueger, G.S. Hotamisligil, R.A. Clark, and T.S. Kupper. 2017. Survival of tissue-resident memory T cells requires exogenous lipid uptake and metabolism. Nature. 543:252–256. doi:10.1038/nature21379.

49. Park, S.L., A. Zaid, J.L. Hor, S.N. Christo, J.E. Prier, B. Davies, Y.O. Alexandre, J.L. Gregory, T.A. Russell, T. Gebhardt, F.R. Carbone, D.C. Tscharke, W.R. Heath, S.N. Mueller, and L.K. MacKay. 2018. Local proliferation maintains a stable pool of tissue-resident memory T cells after antiviral recall responses article. Nat. Immunol. 19:183–191. doi:10.1038/s41590-017-0027-5.

50. Patel, D.F., A. Gaggar, J.E. Blalock, L.G. Gregory, C.M. Lloyd, and R.J. Snelgrove. 2019. An extracellular matrix fragment drives epithelial remodeling and airway hyperresponsiveness. Sci. Transl. Med. 11. doi:10.1126/scitranslmed.aaw0462.

51. Perros, F., H.C. Hoogsteden, A.J. Coyle, B.N. Lambrecht, and H. Hammad. 2009. Blockade of CCR4 in a humanized model of asthma reveals a critical role for DC-derived CCL17 and CCL22 in attracting Th2 cells and inducing airway inflammation. *Allergy Eur*. J. Allergy Clin. Immunol. 64:995–1002. doi:10.1111/j.1398-9995.2009.02095.x.

52. Robinson, M.D., D.J. McCarthy, and G.K. Smyth. 2009. edgeR: A Bioconductor package for differential expression analysis of digital gene expression data. Bioinformatics. 26:139–140. doi:10.1093/bioinformatics/btp616.

53. Schenkel, J.M., K.A. Fraser, V. Vezys, and D. Masopust. 2013. Sensing and alarm function of resident memory CD8 + T cells. Nat. Immunol. 14:509–513. doi:10.1038/ni.2568.

54. Schenkel, J.M., and D. Masopust. 2014. Tissue-resident memory T cells. Immunity. 41:886–897. doi:10.1016/j.immuni.2014.12.007.

55. Shea, B.S., S.F. Brooks, B.A. Fontaine, J. Chun, A.D. Luster, and A.M. Tager. 2010. Prolonged exposure to sphingosine 1-phosphate receptor-1 agonists exacerbates vascular leak, fibrosis, and mortality after lung injury. Am. J. Respir. Cell Mol. Biol. 43:662–673. doi:10.1165/rcmb.2009-0345OC.

56. Shinoda, K., K. Hirahara, T. Iinuma, T. Ichikawa, A.S. Suzuki, K. Sugaya, D.J. Tumes, H. Yamamoto, T. Hara, S. Tani-Ichi, K. Ikuta, Y. Okamoto, and T. Nakayama. 2016. Thy1+ IL-7+ lymphatic endothelial cells in iBALT provide a survival niche for memory T-helper cells in allergic airway inflammation. Proc. Natl. Acad. Sci. U. S. A. 113:E2842–E2851. doi:10.1073/pnas.1512600113.

57. Slütter, B., N. Van Braeckel-Budimir, G. Abboud, S.M. Varga, S. Salek-Ardakani, and J.T. Harty. 2017. Dynamics of influenza-induced lung-resident memory T cells underlie waning heterosubtypic immunity. Sci. Immunol. 2. doi:10.1126/sciimmunol.aag2031.

58. Smolders, J., K.M. Heutinck, N.L. Fransen, E.B.M. Remmerswaal, P. Hombrink, I.J.M. ten Berge, R.A.W. van Lier, I. Huitinga, and J. Hamann. 2018. Tissue-resident memory T cells populate the human brain. Nat. Commun. 9:1–14. doi:10.1038/s41467-018-07053-9.

59. Snelgrove, R.J., L.G. Gregory, T. Peiró, S. Akthar, G.A. Campbell, S.A. Walker, and C.M. Lloyd. 2014. Alternaria-derived serine protease activity drives IL-33-mediated asthma exacerbations. J. Allergy Clin. Immunol. 134:24–26. doi:10.1016/j.jaci.2014.02.002.

60. Soriano, J.B., A.A. Abajobir, K.H. Abate, S.F. Abera, A. Agrawal, M.B. Ahmed, A.N. Aichour, I. Aichour, M.T. Eddine Aichour, K. Alam, N. Alam, J.M. Alkaabi, F. Al-Maskari, N. Alvis-Guzman, A. Amberbir, Y.A. Amoako, M.G. Ansha, J.M. Antó, H. Asayesh, T.M. Atey, E.F.G.A. Avokpaho, A. Barac, S. Basu, N. Bedi, I.M. Bensenor, A. Berhane, A.S. Beyene, Z.A. Bhutta, S. Biryukov, D.J. Boneya, M. Brauer, D.O. Carpenter, D. Casey, D.J. Christopher, L. Dandona, R. Dandona, S.D. Dharmaratne, H.P. Do, F. Fischer, T.T. Gebrehiwot, A. Geleto, A.G. Ghoshal, R.F. Gillum, I.A. Mohamed Ginawi, V. Gupta, S.I. Hay, M.T. Hedayati, N. Horita, H.D. Hosgood, M.M.B. Jakovljevic, S.L. James, J.B. Jonas, A. Kasaeian, Y.S. Khader, I.A. Khalil, E.A. Khan, Y.H. Khang, J. Khubchandani, L.D. Knibbs, S. Kosen, P.A. Koul, G.A. Kumar, C.T. Leshargie, X. Liang, H. Magdy Abd El Razek, A. Majeed, D.C. Malta, T. Manhertz, N. Marquez, A. Mehari, G.A. Mensah, T.R. Miller, K.A. Mohammad, K.E. Mohammed, S. Mohammed, A.H. Mokdad, M. Naghavi, C.T. Nguyen, G. Nguyen, Q. Le Nguyen, T.H. Nguyen, D.N.A. Ningrum, V.M. Nong, J.I. Obi, Y.E. Odeyemi, F.A. Ogbo, E. Oren, P.A. Mahesh, E.K. Park, G.C. Patton, K. Paulson, M. Qorbani, R. Quansah, A. Rafay, M.H.U. Rahman, R.K. Rai, S. Rawaf, M. Reinig, et al. 2017. Global, regional, and national deaths, prevalence, disability-adjusted life years, and years lived with disability for chronic obstructive pulmonary disease and asthma, 1990–2015: a systematic analysis for the Global Burden of Disease Study 2015. Lancet Respir. Med. 5:691–706. doi:10.1016/S2213-2600(17)30293-X.

61. Stary, G., A. Olive, A.F. Radovic-Moreno, D. Gondek, D. Alvarez, P.A. Basto, M. Perro, V.D. Vrbanac, A.M. Tager, J. Shi, J.A. Yethon, O.C. Farokhzad, R. Langer, M.N. Starnbach, and U.H. Von Andrian. 2015. A mucosal vaccine against Chlamydia trachomatis generates two waves of protective memory T cells. Science (80-.). 348. doi:10.1126/science.aaa8205.

62. Strutt, T.M., K. Dhume, C.M. Finn, J.H. Hwang, C. Castonguay, S.L. Swain, and K.K. McKinstry. 2018. IL-15 supports the generation of protective lung-resident memory CD4 T cells. Mucosal Immunol. 11:668–680. doi:10.1038/mi.2017.101.

63. Szabo, P.A., M. Miron, and D.L. Farber. 2019. Location, location, location: Tissue resident memory T cells in mice and humans. Sci. Immunol. 4:1–12. doi:10.1126/sciimmunol.aas9673.

64. Takamura, S., H. Yagi, Y. Hakata, C. Motozono, S.R. McMaster, T. Masumoto, M. Fujisawa, T. Chikaishi, J. Komeda, J. Itoh, M. Umemura, A. Kyusai, M. Tomura, T. Nakayama, D.L. Woodland, J.E. Kohlmeier, and M. Miyazawa. 2016. Specific niches for lung-resident memory CD8+ T cells at the site of tissue regeneration enable CD69-independent maintenance. J. Exp. Med. 213:3057–3073. doi:10.1084/jem.20160938.

65. Tibbitt, C.A., J.M. Stark, L. Martens, J. Ma, J.E. Mold, K. Deswarte, G. Oliynyk, X. Feng, B.N. Lambrecht, P. De Bleser, S. Nylén, H. Hammad, M. Arsenian Henriksson, Y. Saeys, and J.M. Coquet. 2019. Single-Cell RNA Sequencing of the T Helper Cell Response to House Dust Mites Defines a Distinct Gene Expression Signature in Airway Th2 Cells. Immunity. 51:169–184.e5. doi:10.1016/j.immuni.2019.05.014.

66. Turner, D.L., M. Goldklang, F. Cvetkovski, D. Paik, J. Trischler, J. Barahona, M. Cao, R. Dave, N. Tanna, J.M. D’Armiento, and D.L. Farber. 2018. Biased Generation and In Situ Activation of Lung Tissue-Resident Memory CD4 T Cells in the Pathogenesis of Allergic Asthma. J. Immunol. ji1700257. doi:10.4049/jimmunol.1700257.

67. Wakim, L.M., A. Woodward-Davis, R. Liu, Y. Hu, J. Villadangos, G. Smyth, and M.J. Bevan. 2012. The Molecular Signature of Tissue Resident Memory CD8 T Cells Isolated from the Brain. J. Immunol. 189:3462–3471. doi:10.4049/jimmunol.1201305.

68. Walker, J.A., and A.N.J. McKenzie. 2018. TH2 cell development and function. Nat. Rev. Immunol. 18:121–133. doi:10.1038/nri.2017.118.

69. Wolf, A.M., K. Eller, R. Zeiser, C. Dürr, U. V. Gerlach, M. Sixt, L. Markut, G. Gastl, A.R. Rosenkranz, and D. Wolf. 2009. The Sphingosine 1-Phosphate Receptor Agonist FTY720 Potently Inhibits Regulatory T Cell Proliferation In Vitro and In Vivo. J. Immunol. 183:3751–3760. doi:10.4049/jimmunol.0901011.

70. Woodruff, P.G., B. Modrek, D.F. Choy, G. Jia, A.R. Abbas, A. Ellwanger, J.R. Arron, L.L. Koth, and J. V. Fahy. 2009. T-helper type 2-driven inflammation defines major subphenotypes of asthma. Am. J. Respir. Crit. Care Med. 180:388–395. doi:10.1164/rccm.200903-0392OC.

